# Integrated temporal profiling of iPSCs-derived motor neurons from ALS patients carrying *C9orf72*, *FUS*, *TARDBP*, and *SOD1* mutations

**DOI:** 10.1101/2024.10.15.618602

**Authors:** Guo-ming Ma, Cong-cong Xia, Bo-yu Lyu, Jie Liu, Fang Luo, Ming-feng Guan, Jun-ying Wang, Li Sun, Lin Zhang, Yan Chen, Ying-wei Mao, Guo-qiang Yu, Wen-yuan Wang

**Affiliations:** Interdisciplinary Research Center on Biology and Chemistry, Shanghai Institute of Organic Chemistry, Chinese Academy of Science, Shanghai, 200032, China; Bradley Department of Electrical and Computer Engineering, Virginia Polytechnic Institute and State University, Arlington, VA 22203, USA; The International Peace Maternity and Child Health Hospital, School of Medicine, Shanghai Jiao Tong University, Shanghai 200030, China; Research Center for Aging and Medicine, Huashan Hospital, Fudan University, 12 Wulumuqi Zhong Road, Jing’an District, Shanghai, 200040, China; Department of Neurology & National Clinical Research Center for Aging and Medicine, Huashan Hospital, Fudan University, 12 Wulumuqi Zhong Road, Shanghai, 200040, China; 214 Life Sciences Building, Penn State University, University Park, PA 16802, USA; Department of Rehabilitation Medicine, Huashan Hospital, Fudan University, Shanghai, 200040, China

**Keywords:** Amyotrophic lateral sclerosis (ALS), Induced pluripotent stem cells (iPSC), Motor neurons (MN), RNA sequencing (RNA-Seq)

## Abstract

Amyotrophic Lateral Sclerosis (ALS) is a lethal neurodegenerative disease that damages motor neurons in the central nervous system, causing progressive muscle weakness that ultimately leads to death. However, its underlying mechanisms still need to be fully understood, particularly the heterogeneity and similarity between various gene mutants during disease progression. In this study, we conducted temporal RNA-seq profiling in human induced pluripotent stem cells (hiPSCs) and iPSC-derived motor neurons (iMNs) carrying the *C9orf72*, *FUS*, *TARDBP*, and *SOD1* mutations from both ALS patients and healthy individuals. We discovered dysregulated gene expression and alternative splicing (AS) throughout iMN development and maturation, and ALS iMNs display enrichment of cytoskeletal defects and synaptic alterations from premature stage to mature iMNs. Our findings indicate that synaptic gene dysfunction is the common molecular hallmark of fALS, which might result in neuronal susceptibility and progressive motor neuron degeneration. Analysis of upstream splicing factors revealed that differentially expressed RNA-binding proteins (RBPs) in ALS iMNs may cause abnormal AS events, suggesting the importance of studying RBP defects in ALS research. Overall, our research provides a comprehensive and valuable resource for gaining insights into the shared mechanisms of ALS pathogenesis during motor neuron development and maturation in iMN models.

## INTRODUCTION

Amyotrophic Lateral Sclerosis (ALS) is a fatal neurodegenerative disorder that causes progressive weakness and muscle atrophy due to the loss of motor neurons in the spinal cord and brain^1,2^. It has a median survival of 3 to 5 years after diagnosis, and there is currently no effective treatment available^1,2^. Around 10-15% of patients have a positive family history of ALS, and 70% of familial cases have mutations within known ALS genes^3^. Over 40 genes have been associated with ALS, with most cases linked to *C9orf72*, *SOD1*, *TARDBP*, and *FUS* mutations, although the frequency of genetic subtypes varies by population ancestry^3–5^. The mechanisms behind ALS are still unclear but broadly fall into impaired RNA metabolism, altered proteostasis or autophagy, cytoskeletal or trafficking defects, and mitochondrial dysfunction^6,7^. Aggregates of RNA-binding protein transactive response (TAR) DNA-binding protein 43 (TDP-43) and fused in sarcoma/translocated in liposarcoma (FUS/TLS) have been observed in FTD and ALS patients’ brains and spinal cords^8–15^. It is emerging that TDP-43 and FUS mutations in both proteins provoke axonal translation defects before disease development, suggesting that dysfunctional local translation may contribute to neurodegeneration^16–18^. ALS’s most common genetic determinant is a hexanucleotide repeat expansion in the chromosome 9 open reading frame 72 (*C9orf72*) gene^5,19–21^. Possible pathogenic mechanisms underlying the *C9orf72* repeat expansion have been attributed to both gain-of-function (GOF) and loss-of-function (LOF) effects^22–29^. Superoxide dismutase 1 (*SOD1*) is the first identified ALS gene in 1993^30,31^. Many SOD1 variants have been described in ALS patients, and studies identify mutant SOD1 secretion as a double-edged sword in ALS pathogenesis^5,22,32–37^. However, the relationship between the genetic defect and the pathophysiology of the disease remains unclear.

Although the genes responsible for causing ALS may appear to be unrelated, recent studies have demonstrated that similar critical cellular pathways exhibit abnormalities in ALS patients with various genetic backgrounds. These pathways include unfolded protein response, DNA damage response, mitochondrial damage, excitotoxicity, axonal transport defects, dysregulation of RNA metabolism, and endoplasmic reticulum (ER) stress^1,3^. These common molecular mechanisms suggest that different ALS genes share many similarities in causing the disease. These results reveal the complexity of the pathogenic mechanisms of ALS.

The use of induced pluripotent stem cell (iPSC)-derived motor neuron (iMNs) models has dramatically expanded our ability to model ALS with its clinical and genetic diversity^38–41^. Biobanks such as NeuroLINCS (http://neurolincs.org/) and Answer ALS have increased the scope of ALS iPSC-derived motor neuron (iMN) research, providing an opportunity to identify common motor neuron abnormalities across different ALS genetic backgrounds^42^.

Given the diversity and complexity of ALS pathogenesis, a crucial question arises: What molecular mechanisms, common or specific, are involved in ALS with different causative genes? Many attempts to figure out the pathological mechanisms underlying ALS have initially focused on single ALS genes, which are inadequate to elucidate ALS pathogenesis fully. The onset of ALS commonly occurs in mid-adulthood, at around 55 years of age; however, the effects of ALS genes may manifest earlier in life^28^. Hence, we aimed to elucidate molecular level changes throughout motor neuron development and systematically examine the effects of ALS-causing genes on transcriptomes during motor neuron differentiation. iMN is perfectly positioned to explore the functional consequences of genetic variants on motor neurons during the initial phases of the disease.

In this study, we generated fibroblast cell lines from ALS patients bearing *C9orf72*, *FUS*, *TARDBP*, and *SOD1* mutations, respectively, reprogrammed the cells into iPSCs, and differentiated them into motor neurons to investigate the pathogenesis of ALS in vitro. We addressed the common and specific gene expression changes, alternative splicing (AS) dysregulation, and specific splicing factors regulating AS events throughout motor neuron development and ALS progression in *C9orf72*-ALS, *FUS*-ALS, *TARDBP*-ALS, and *SOD1*-ALS by using a combination of iPSC differentiation technology and temporal RNA-seq profiling.

Our analysis highlighted the synaptic function-related genes and pathways underlying shared mechanisms, regulated by a broad range of ALS disease risk factors, that lead to neuronal vulnerability and progressive degeneration. These results suggested that the effects of ALS genes on transcriptome changes are evident in the early stages of motor neuron differentiation. Our findings provide a valuable resource for the ALS research community.

## Results

### Generation of iPSCs and differentiation of functional iMNs from ALS patients

To examine the common effect of mutations of ALS causative genes on iMNs, we generated iPSCs from nine fibroblasts of ALS patients carrying *SOD1* (*SOD1*-1, *SOD1*-2, *SOD1*-3, and *SOD1*-4), *TARDBP* (*TARDBP*-1 and *TARDBP*-2), *FUS* (*FUS*-1 and *FUS*-2), and *C9orf72* gene mutations, and four fibroblasts from healthy individuals (Table S1). The karyotype and mutations of all cell lines were experimentally confirmed (Supplementary Fig. 1A, B). We obtained differentiated spinal iMNs through a modified protocol (Fig. 1A), which resulted in a highly near-pure population of PAX6+ and NESTIN+ neuro precursors cells (NPCs) (Supplementary Fig. 2A) in 6 days, OLIG2+ motor neuron progenitors (MNPs) (> 98%, Supplementary Fig. 2B) in 12 days, ISL1+ and HB9+ iMNs (> 80%, Supplementary Fig. 2C, D) in 23 days, and CHAT+ maturing iMNs in 28 days^40^ (Supplementary Fig. 2E). The efficiency of iMN differentiation was similar between control and ALS subgroups.

**Figure 1.**
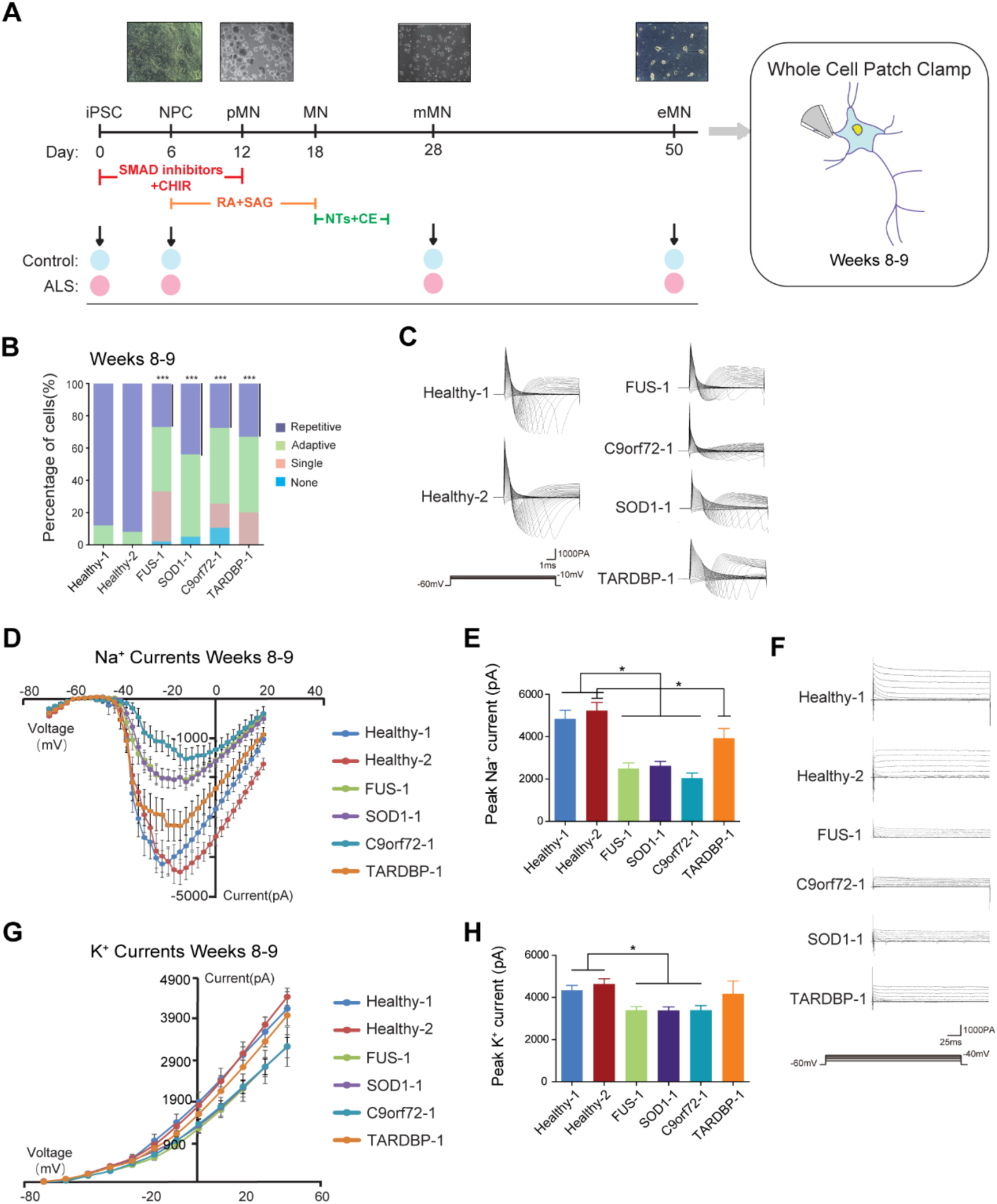
Generation of iPSCs and differentiation of functional iMNs from ALS patients. (A) Schematic of the differentiation process of induced pluripotent stem cells (iPSC) into motor neurons (mMN) from day 0 to day 50. The process involves the transition from induced-pluripotent stem cells (iPSC) to neural progenitor cells (NPC), then to patterned precursor motor neurons (pMN), followed by post-mitotic but electrophysiologically immature motor neurons (yMN) at day 28, and finally resulting in electrophysiologically mature motor neurons (mMN) at day 50. (B) Illustrations of the four distinct categories of action potential (AP) firing observed in iMNs. (C) Proportion of cells in each AP firing category in iMNs from Healthy-1 (n = 58), Healthy-2 (n = 61), *FUS*-1 (n = 52), *SOD1*-1 (n = 41), *C9orf72*-1 (n = 47) and *TARDBP*-1 (n = 41) lines at weeks 8-9 post-plating (student’s t-test; *** p < 0.001). (D) Raw data of fast, inactivating Na^+^ currents from iMNs at weeks 8-9. (E) The current-voltage relationships of peak Na^+^ currents were recorded from iMNs at weeks 8-9. (F) Peak fast, inactivating Na^+^ currents plotted from iMNs at weeks 8-9 (student’s t-test; * p < 0.05; mean ± SEM). (G) Raw data of persistent K^+^ currents from iMNs at weeks 8-9. (H) The current-voltage relationships of peak K^+^ currents were recorded from iMNs at weeks 8-9. (I) Peak K^+^ currents plotted from iMNs at weeks 8-9 (student’s t-test; * p < 0.05; mean ± SEM).

We next sought to evaluate whether iMNs were functional. After co-culturing the iMNs with differentiated mouse C2C12 cells for 7 days, we observed the formation of neuromuscular junctions, as indicated by the presence of aggregated α-BTX+ acetylcholine receptors on myotubes and their overlap with CHAT+ neurites (Supplementary Fig. 2F).

Then, we performed whole-cell patch clamp recordings to access the electrophysiological properties of iMNs from healthy individuals. We found that healthy iMNs were electrophysiologically active, as evidenced by their ability to elicit short-lasting action potentials in response to depolarizing current injection in current-clamp recording (Supplementary Fig. 2G), suggesting they are fully functional iMNs. Taken together, we have successfully developed an iPSC-based human spinal motor neuron disease model of ALS.

### Functional perturbation of ALS patient-derived iMNs

We then investigated whether these ALS iMNs exhibited functional perturbations. To uncover signs of neuronal dysfunction, we examined electrophysiological properties of iMNs by whole-cell patch-clamp recordings (Fig. 1A). Our recordings showed four different firing patterns in response to current injections, including no firing, single, adaptive and repetitive firing^43^. The repetitive firing was characterized by a train of action potentials (APs) lasting for the duration of the square current injection (1s); in contrast, the adaptive firing comprised multiple action potentials that ceased before the end of the current stimuli^43^. Cells were categorized as adaptive if they could not repetitively fire in response to the series of applied current steps^43^. We found that the number of cells able to fire APs showed a significant decrease in ALS iMNs compared with controls at weeks 8-9 (Fig. 1B), especially, the cells that can fire APs repetitively, indicating a reduction in the patient-iMN’s ability to sustain a high rate of electrical activity over time, and the impairment of its physiological functions.

To better understand the reason behind the decrease in the output of AP in ALS iMNs, we conducted tests on voltage-activated currents during AP generation. Our investigation began with fast, inactivating Na^+^ currents, which are responsible for the upstroke of the action potential. We applied a range of voltage steps ranging from −70 to 20 mV in 2.5 mV increments. These voltage steps lasted for 10 ms and were applied from a holding potential of −60 mV (Fig. 1C). Our results showed that there was a gradual decline of Na^+^ currents in ALS iMNs (Fig. 1D). Peak Na^+^ currents significantly decreased in *FUS*, *SOD1* and *C9orf72* iMNs compared to controls groups at weeks 8-9. A slightly decreasing trend was observed in *TARDBP* iMNs (Fig. 1E).

After identifying the progressive Na^+^ currents loss, we investigated whether these results indicated a more general reduction in voltage-activated currents in ALS iMNs. We measured persistent K^+^ currents using a range of voltage steps (−70 to 40 mV, 10 mV increments, 500 ms duration) from a holding potential of −60 mV (Fig. 1F). Our findings showed a progressive loss in peak K^+^ currents in ALS iMNs at weeks 8-9 (Fig. 1G). iMNs harboring *SOD1*, *FUS*, and *C9orf72* mutations exhibited significantly lower K^+^ currents than controls. iMNs derived from *TARDBP*-ALS patients showed a slight reduction in K^+^ currents compared to controls (Fig. 1H). Our data demonstrate a progressive loss in both fast, inactivating Na^+^ currents and persistent, voltage-activated K^+^ currents in elder iMNs from ALS patients. It is plausible that loss of action potential output and the reduction in voltage-activated currents underlie the progressive functional decline in ALS iMNs.

### Transcriptomic disturbances during iMN development in ALS genetic background

To investigate ALS-related transcriptomic changes during human motor neuron development and maturation, we carried out deep high-throughput RNA-sequencing (RNA-seq) from iPSCs, NPCs, young iMNs, and mature iMNs derived from nine patients with the ALS-causing gene mutations and four healthy donors. Hierarchical clustering reveals that gene expression changes were primarily regulated by developmental stage within the motor neuron lineage rather than by genetic background (Fig. 2A). By clustering gene expression patterns based on known gene markers of iPSC, NPC, and motor neuron, we identified critical iPSC-specific gene markers, and significantly up-regulated NPC markers. More importantly, 16 key genes were highly expressed in iMNs on day 28 and day 50, indicating spinal motor neuron (spMN) maturation (Fig. 2B). These results suggest that mutated ALS genes did not disrupt motor neuron maturation; the iMNs can recapitulate in vivo adult motor neuron development and maturation and thus could be an ideal model for studying the progression of ALS pathogenesis.

**Figure 2.**
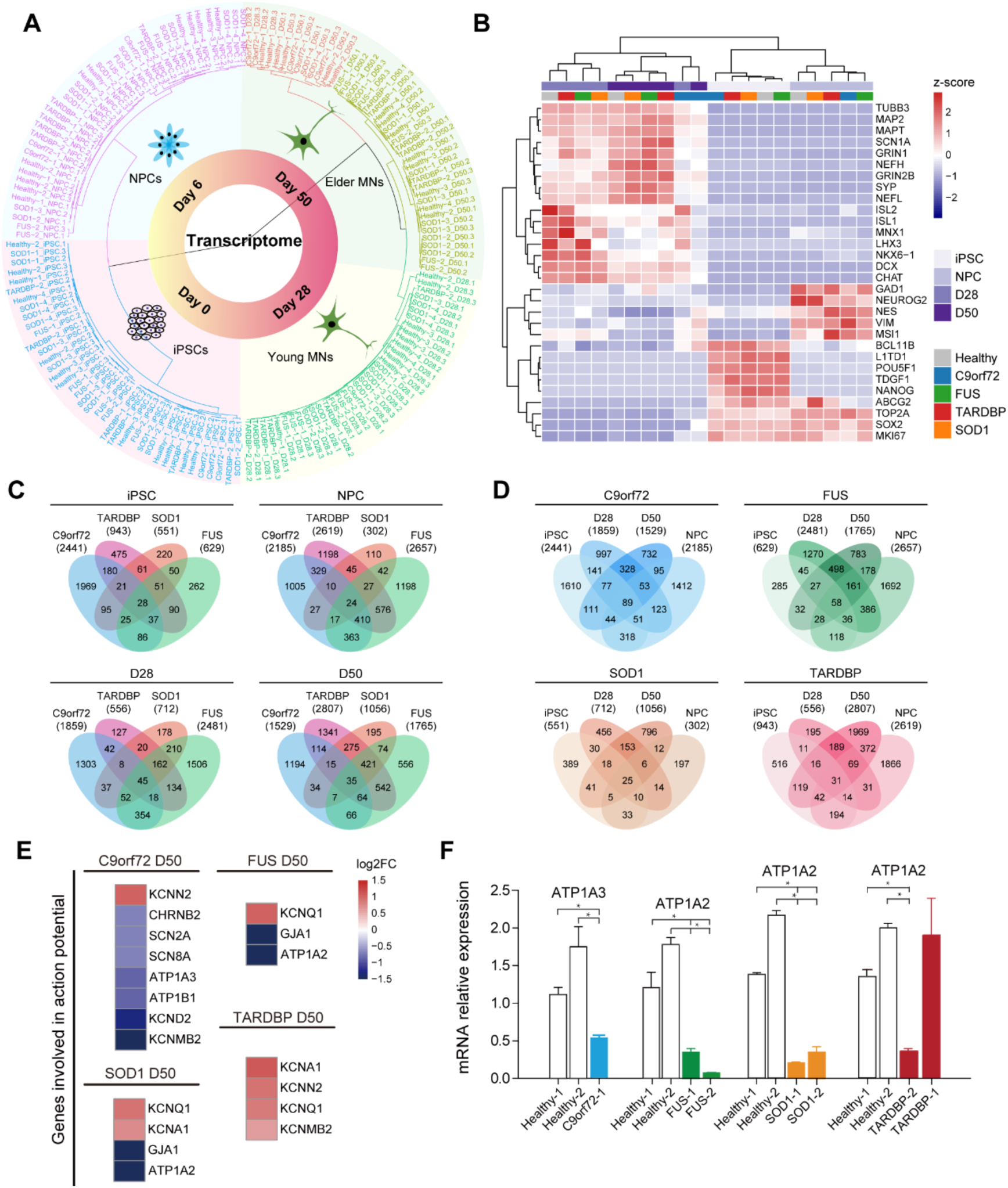
Transcriptomic disturbances during iPSC-derived motor neurogenesis in ALS genetic background. (A) Hierarchical clustering (Manhattan distance and Ward’s method) of all samples by transcriptomes from iPSCs to iMNs. (B) Heatmap of the normalized average expression of selected marker genes related to pluripotency, NPCs and spinal MNs (spMNs) maturation across iMN developmental stages. (C) Overlap of differentially expressed genes (DEGs) of ALS subgroups compared with healthy controls for each stage (FDR < 0.05, |fold change| ≥ 1.5). (D) Overlap of DEGs across four iMN developmental stages in each ALS subgroup (FDR < 0.05, |fold change| ≥ 1.5). (E) Heatmap of genes related to generating AP showing expression changes in mature iMNs on day 50 in each ALS subgroup. (F) Bar plot showing expression changes of genes related to generating APs on day 50 measured by qPCR (student’s t-test; * p < 0.05, ** p < 0.01, *** p < 0.001, **** p < 0.0001; mean ± SEM).

We then investigated how ALS genes commonly and differentially affect changes in gene expression at the transcriptome level in human motor neurons. We found that certain genes were expressed differently in ALS subgroups than in healthy subgroups at each stage of motor neuron development, which were referred to as differentially expressed genes (DEGs; |fold change| ≥ 1.5, FDR < 0.05). By comparing the DEGs of various ALS genes at each time point, we discovered that a significant number of genes showed altered expression levels in various ALS subgroups (Fig. 2C). Furthermore, we found that many genes showed altered expression patterns throughout the early developmental stages of iMNs, which persisted into later stages, the progressive disease state, caused by ALS genes (Fig. 2D). Therefore, it is important to elucidate the molecular mechanisms that are commonly and temporally impacted by *SOD1*, *FUS*, *TARDBP*, and *C9orf72* mutations to gain a better understanding of ALS pathogenesis.

To decipher the root cause of the abnormal electrophysiological properties observed earlier, we analyzed gene expression changes on day 50. The results revealed significant expression changes of many genes involved in generating action potentials in ALS iMNs (Fig. 2D), particularly subunits of voltage-gated sodium channels (such as *SCN2A*), subunits of the voltage-gated potassium channels (e.g., *KCNQ1*, and *KCNA1*) and catalytic subunits of Na^+^/K^+^-ATPase (e.g. *ATP1A3* and *ATP1A2*). These genes play crucial roles in establishing and maintaining the electrochemical gradients of Na^+^ and K^+^ across the plasma membrane. Notably, *ATP1A2* was down-regulated in both *FUS* and *SOD1* iMNs, and *ATP1A3* was down-regulated in *C9orf72* iMNs (Fig. 2E). The dysregulation of these genes was further validation by quantitative PCR (Fig. 2F). Intriguingly, *ATP1A2* was also down-regulated in *TARDBP*-2 lines but not in *TARDBP*-1 lines, in line with electrophysiological data showing no significant differences between iMNs from *TARDBP*-2 lines and healthy lines. These results strongly suggest that the downregulation of Na^+^/K^+^-ATPases is a common molecular mechanism underlying ALS-associated neuronal dysfunction caused by ALS genes, thereby directly linking our findings to the disease state.

### ALS iMNs exhibit dysregulated neuronal functions on gene expression level across various genetic backgrounds

There are currently more than 100 genes are associated with ALS patients, of them, 39 have a strong correlation and are considered to be causative, while others are linked to or modify disease initiation and progression by increasing the likelihood of developing ALS. Although these genes may seem individually unrelated, some are involved in the same cellular pathways, such as axonal transport defects, unfolded protein response, and ER stress, suggesting that these cellular events play a crucial role in the proper function of motor neuron circuitry.

To investigate possible connections among different ALS genes involved in the pathogenesis of ALS, we compared the levels of gene expression in day 50 iMNs from ALS patients versus healthy individuals. We identified unique DEGs that were implicated in each ALS subgroup. Using Gene Ontology (GO) enrichment analysis, we found ten significantly enriched biological processes (BP) for DEGs in ALS-*C9orf72*, ALS-*FUS*, ALS-*TARDBP*, and ALS-*SOD1*, which we presented in Fig. 3 (adjusted p-value < 0.05). We found that ALS-*C9orf72* had affected synaptic transmission and ion transport (Fig. 3A). iMNs with ALS-*FUS* showed extracellular matrix organization, synaptic transmission, and DNA damage (Fig. 3B). ALS-*TARDBP* had affected extracellular matrix organization, regulation of neuronal differentiation, and nonsense-mediated decay (Fig. 3C). iMNs with ALS-*SOD1* had changes in extracellular matrix organization, DNA damage, and oxidative stress response (Fig. 3D). Our comparisons of significant BPs from these four ALS genes identified several common dysregulated cellular functions. From the NPCs to the D50-iMNs, cytoskeleton, cell adhesion, cellular composition organization, synaptic function, cellular response, and neuronal development were enriched in two or more ALS subgroups, indicating that common dysregulated neuronal mechanisms may be involved among these ALS-causative genes (Fig. 3E). These results suggest that alterations on transcriptome level are usually distinct in iMNs derived from ALS patients with mutations in *C9orf72, FUS*, *TARDBP* and *SOD1*, but may overlap to some extent in mature iMNs of disease onset.

**Figure 3.**
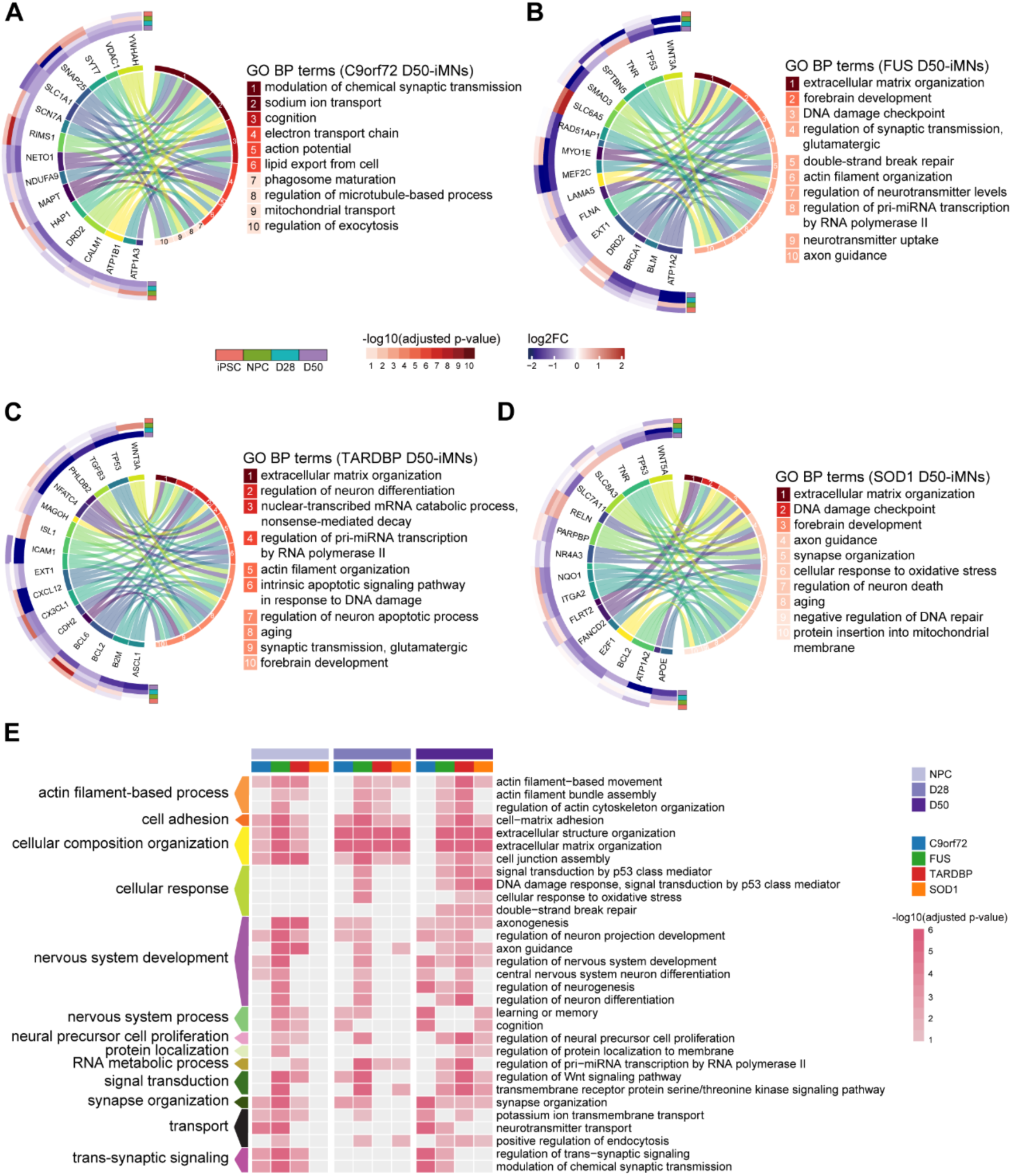
Distinct and common transcriptome alterations in mature iMNs across four ALS subgroups. (A-D) Circos plot showing significantly dysregulated genes and gene ontology (GO) enrichment in ALS-*C9orf72* (A), ALS-*FUS* (B), ALS-*TARDBP* (C), and ALS-*SOD1* (D) iMNs on day 50, respectively (adjusted p-value < 0.05). (E) Heatmap showing the enrichment of selected GO terms resulting from DEGs in each ALS subgroup from NPC to D50-iMN stages (adjusted p-value < 0.05).

### ALS genes affect temporal dynamics of gene expression and exhibits dysregulated neuronal functions from premature iMN stages

We then examined the time-dependent gene expression changes in ALS subgroups. We clustered the log2 fold change values of DEGs across all stages of development and performed GO analysis of DEGs from each cluster. This analysis helped us identify the effect of ALS genes on specific patterns of gene expression changes. Our analysis revealed that in ALS-*C9orf72*, cluster 1 displayed a down-regulated pattern from day 28. The genes in this cluster were significantly enriched in the synaptic area mitochondrial function (Fig. 4A). DEGs in cluster 3 were mainly downregulated from the NPC stage, with significant localization in the distal axon and pre-synapse (Fig. 4A). These findings suggest that *C9orf72* mutations may cause abnormalities in synapse and mitochondria from early stages, persisting into elder mature motor neurons. In *TARDBP* iMNs, DEGs in clusters 1, 2 and 3 were significantly down-regulated on day 50, and these genes already showed a trend of down-regulated expression from day 28 (Fig. 4B). These DEGs were enriched in RNA metabolism, protein localization, and DNA damage response. In *FUS* iMNs, clusters 1, 2 and 3 showed a down-regulation trend from day 28. *FUS* mutations lead to functional disorders such as p53 signal transduction and cytoskeleton. (Fig. 4C). These results suggest that *TARDBP* and *FUS* mutations can lead to dysregulation of p53 signal transduction from the early stages of iMN maturation. In *SOD1* iMNs, clusters 1 and 2 represented down-regulated DEGs from day 28; they were enriched in the transforming the growth factor beta receptor signaling pathway and DNA damage response (Fig. 4D). On the other hand, DEGs in cluster 3 showed an up-regulated trend from the NPC stage. They are enriched in synapse organization, postsynaptic endocytosis, learning and memory (Fig. 4D). These results suggest that *SOD1* mutations may cause abnormal synaptic function in the early stages. Overall, our findings reveal that ALS genes can affect many essential motor neuron functions from early stages, which might provide potential early targets for clinical intervention for ALS disease.

**Figure 4.**
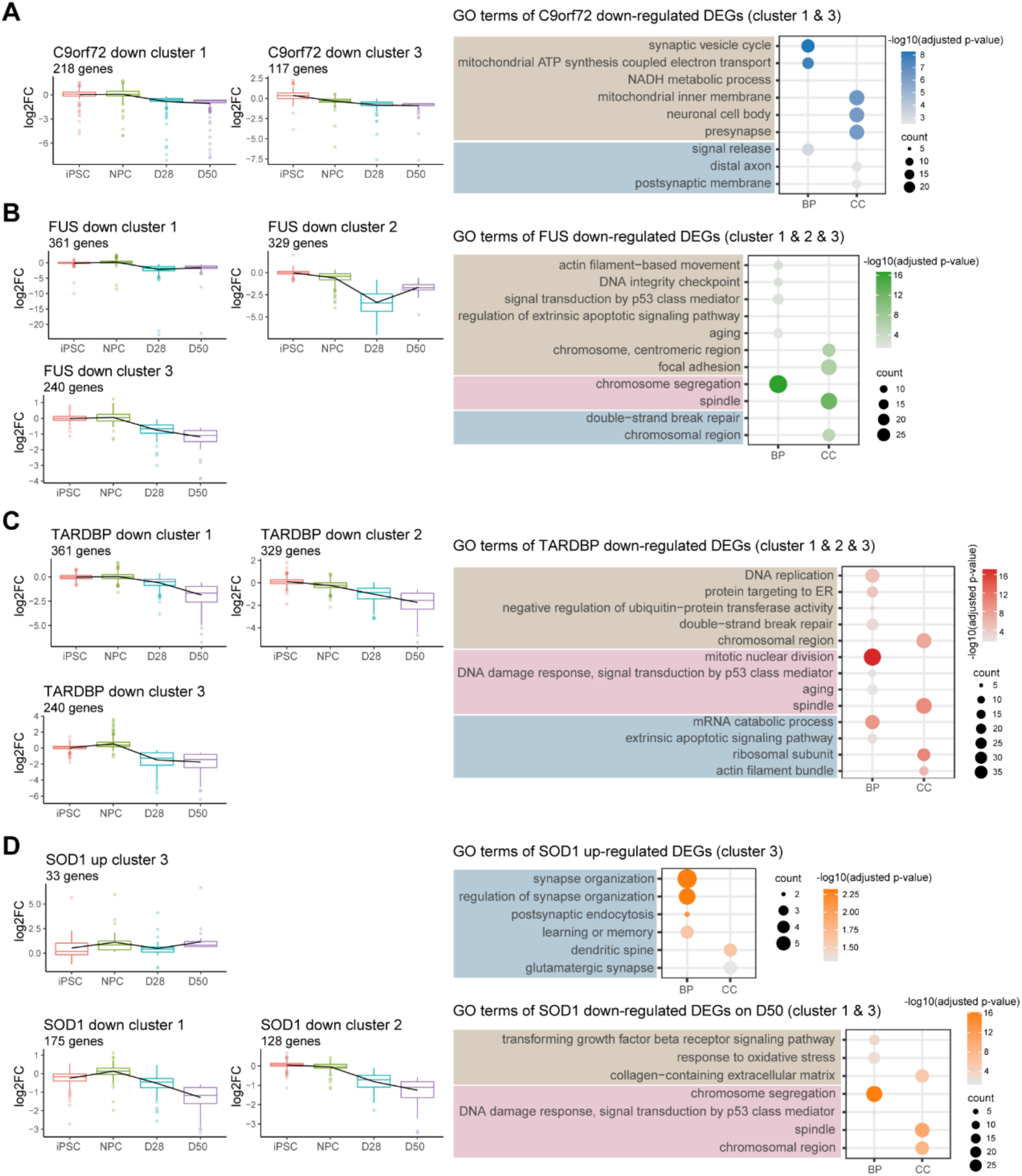
Temporal transcriptome alterations in ALS subgroups during iMN development and maturation. (A-D) Left, trends of log2 fold change values of DEGs from the selected clusters in ALS-*C9orf72* (A), ALS-*FUS* (B), ALS-*TARDBP* (C), and ALS-*SOD1* (D). Right, enriched GO terms in corresponding clusters in ALS-*C9orf72* (A), ALS-*FUS* (B), ALS-*TARDBP* (C), and ALS-*SOD1* (D) (adjusted p-value < 0.05).

### Transcriptome alterations show intricate dynamics of signaling pathway in ALS iMNs especially during mature motor neuron stages

We analyzed the gene expression changes during iMN development and identified DEGs between two consecutive developmental stages to understand the temporal dynamics of the process. Interestingly, we found that *FUS*, *TARDBP*, and *SOD1* mutations had more consistent effects on gene expression levels (Fig. 5A-C). We also performed a Signaling Pathway RespOnsive GENes (PROGENy) analysis to investigate the activated pathways during iMN development and maturation^44^. Our findings revealed that WNT pathway activity increased in healthy and ALS subgroups. However, the JAK-STAT pathway showed decreased activity from iPSCs to NPCs (Fig. 5D), and the EGFR and MAPK pathways were also observed to have decreased activity across all subgroups during the transition from NPCs to mature iMNs (Fig. 5E).

**Figure 5.**
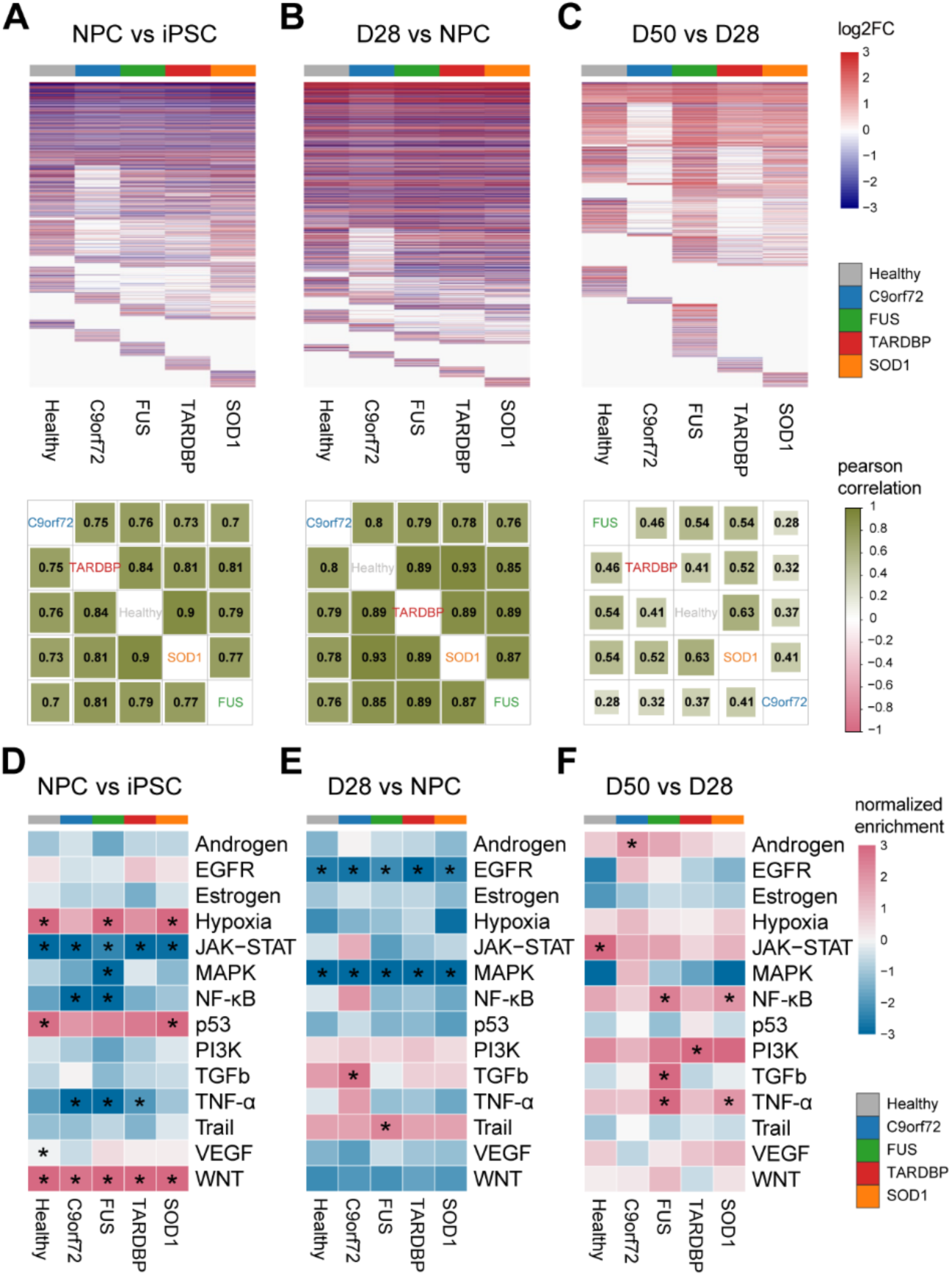
Temporal profiles of transcriptome alterations between each stage of iMN development and maturation. (A-C) Upper, heatmaps of genes differentially expressed in at least one genetic background between two consecutive developmental stages. Lower, the Pearson’s correlation coefficient for transcriptome-wide changes between two genetic backgrounds in NPCs compared to iPSCs (A), D28-iMNs compared to NPCs (B), and D50-iMNs compared to D28-iMNs (C). (D-F) PROGENy signaling pathway activities in NPCs compared to iPSCs (D), D28-iMNs compared to NPCs (E), and D50-iMNs compared to D28-iMNs (F) in healthy controls and ALS subgroups. Pathways that increase in later stages are red, while pathways that decrease are blue. Statistics are from the weighted mean method (enrichment test; * p < 0.05).

At the mature iMN stage, our research uncovered significant differences in the activities of signaling pathways among healthy and various ALS subgroups. Specifically, we observed that JAK-STAT pathway activation was present only in healthy iMNs, while inflammation-related NF-κB and TNF-α pathways were significantly activated in *FUS* and *SOD1* iMNs (Fig. 5F; Supplementary Fig. 3A-D). These findings underscore the substantial divergent expression changes among ALS subgroups at transitioning from D28-iMNs to D50-iMNs, highlighting the divergent pathway activities between healthy and ALS iMNs.

### Healthy and ALS-specific AS events suggests affected neuronal functions in ALS iMNs during iMN differentiation

The precise implementation of specific alternative splicing (AS) processes is essential for brain development and maintaining cell homeostasis^45,46^. In patients with ALS, abnormal AS changes occur in the primary motor cortex, which may lead to protein dysfunction and worsen the progression of the disease^47,48^. To investigate how AS is affected in ALS iMNs during their development and maturation, we conducted AS analyses using rMATS on healthy and ALS subgroups at different stages (Fig. 6A). our analysis identified significant AS events (FDR < 0.05; |InclusionLevelDifference| > 0.1), including skipped exons (SEs), mutually exclusive exons (MXEs), alternative 5’/3’ splice sites (A5SSs, A3SSs), and retained introns (RIs). GO analysis revealed that ALS-specific and healthy-specific alternative spliced genes were enriched in regulating GTPase activity, dendritic or neuron projection, microtubule, and cell polarity from NPC to D28 (as shown in Supplementary Fig. 4B).

**Figure 6.**
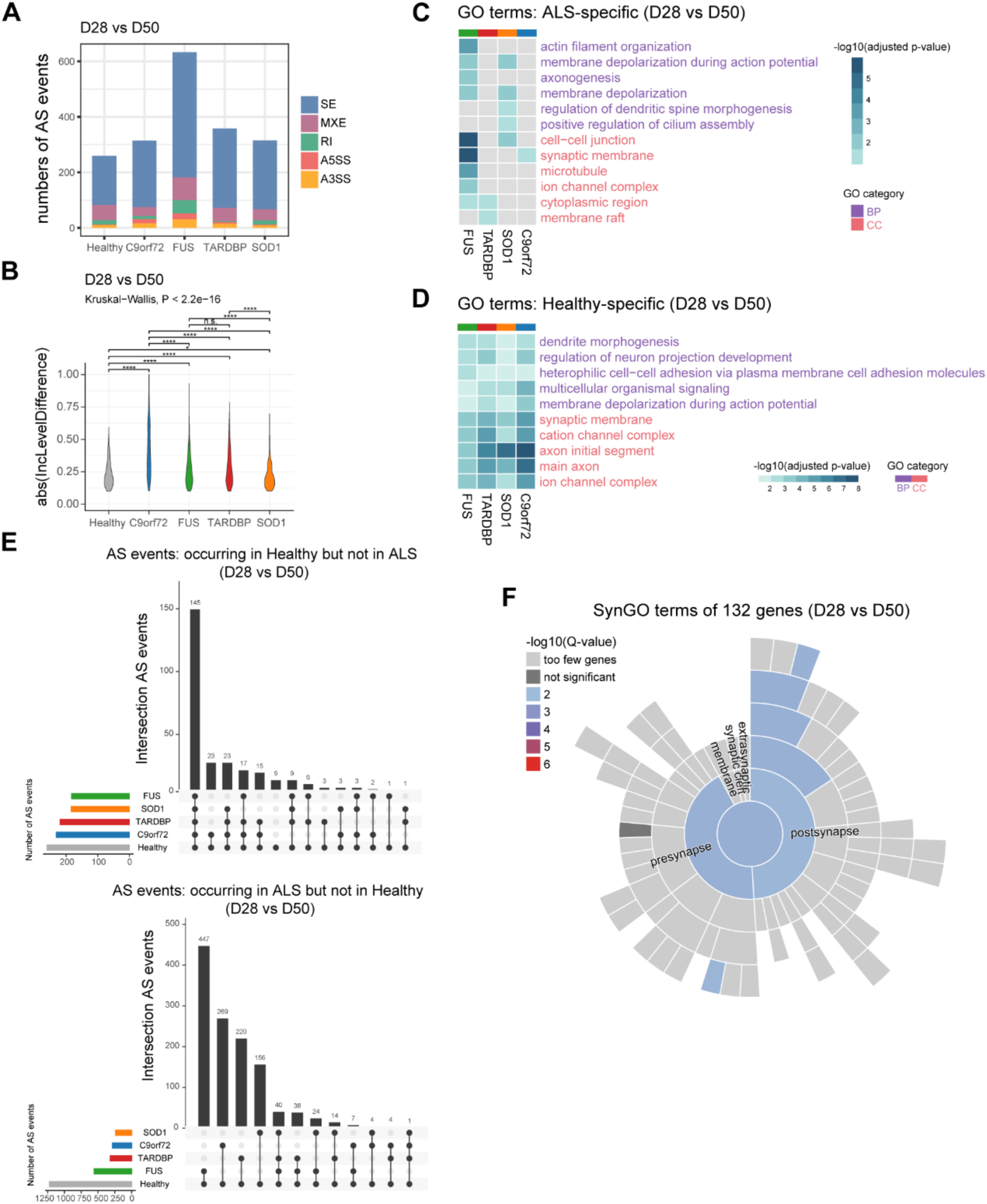
Aberrant splicing events in D50-iMNs compared to D28-iMNs. (A) Numbers of splicing types in ALS subgroups and healthy controls (FDR < 0.05, |InclusionLevelDifference| > 0.1). skipped exon (SE); mutually exclusive exons (MXE); Retained introns (RI); alternative 5’ splice site (A5SS); alternative 3’ splice site (A3SS). (B) Violin plot of the absolute inclusion level differences of significant AS events in ALS subgroups and healthy controls (Wilcoxon signed-rank test; * p < 0.05, ** p < 0.01, *** p < 0.001, **** p < 0.0001, n.s., not significant). (C) Compared to healthy controls, the common GO terms associated with genes only differentially alternative spliced in ALS iMNs from day 28 to day 50 transition stage. (D) Common GO terms associated with genes only differentially alternative spliced in healthy iMNs from day 28 to day 50 transition stage. (E) Overlaps of splicing events significantly altered in healthy controls but not altered in ALS subgroups (left) and vice versa (right). (F) The Sunburst plot displays SynGO annotations for genes with AS in healthy controls but does not show same splicing events in ALS subgroups relative to a background set of brain-expressed genes.

Since ALS typically manifests functional deficits in motor neurons during the later stages of disease progression, we focused on AS events from D28-iMNs to D50-iMNs, and identified 633, 358, 315, 314, and 260 significant AS events in *FUS*, *TARDBP*, *SOD1*, *C9orf72* and healthy subgroups, respectively (Fig. 6A; Supplementary Fig. 4A). Both ALS and healthy subgroups exhibited an increase in the proportion of SE events frequency of iMN terminal maturation (Supplementary Fig. 4A). Notably, it was found that *FUS* iMNs exhibited more splicing events changes than others (Fig. 6A). Additionally, compared to healthy controls, *FUS*, *TARDBP* and *C9orf72* iMNs displayed significantly higher differences in the absolute inclusion level of AS events (Fig. 6B).

We analyzed the inclusion events that were significantly changed in ALS and healthy subgroups (Supplementary Fig. 4C). We identified 556, 317, 239, and 283 AS events that were significant in *FUS*, *TARDBP*, *SOD1*, and *C9orf72* iMNs, respectively. These events were not significant in healthy subgroups. In contrast, we found 183, 219, 184, and 231 AS events to be healthy-specific. Mutant-specific AS events were enriched in GO categories such as actin filament organization and axongenesis in *FUS* iMNs, regulation of dendritic spine morphogenesis in *SOD1* iMNs, and membrane depolarization in *FUS* and *SOD1* iMNs (Fig. 6C). Additionally, we also found those genes undergone alternative splicing are also present in various cellular components, such as synaptic membrane in *FUS* and *C9orf72* iMNs (Fig. 6C).

The genes that display AS events specific to healthy subgroups show more significant functional enrichment. For example, we observed biological processes related to the regulation of neuron projection development, membrane depolarization during action potentials, and cellular components related to the synaptic membrane, axon initial segment and ion channel complex (Fig. 6D). We also focused on genes with differential splicing between healthy and ALS subgroups and found 145 AS events occurring in healthy subgroups but not in ALS subgroups from day 28 to day 50 (Fig. 6E). Interestingly, iMNs with *FUS*, *TARDBP* or *SOD1* mutations showed similar exon inclusion level difference on day 50 (Supplementary Fig. 4D, 4E).

Since aberrant AS events were synaptically enriched in each ALS subgroup respectively, we used SynGO to investigate 132 genes showing different splicing situations of 145 events^49^ (Fig. 6E). Synaptic ontology terms demonstrated enrichment in these genes, including *NRXN2*, *SCN2A*, *LRRC7* and others (Fig. 6F). In conclusion, our study indicates that mutations in ALS causative genes can lead to aberrant alternative splicing, particularly affecting genes specifically spliced in healthy subgroups, which in turn, may impact synaptic and other neuronal functions in motor neuron maturation from day 28 to day 50.

### Aberrant alternative splicing events presents in mature ALS iMNs

We have found that ALS iMNs exhibit abnormal splicing events during development, leading to synaptic dysfunction (Fig. 6). To understand the impact of disease mutations of *FUS*, *TARDBP*, *SOD1*, and *C9orf72* on RNA splicing during disease progression, we analyzed AS events between ALS and healthy subgroups (Fig. 7A). We noticed significant changes in AS from NPCs to D50-iMNs, with SE events accounting for approximately 60% of the AS events, which is consistent with previous studies^50^ (Fig. 7B; Supplementary Fig. 5A). AS changes at NPC stage were more significant than mature iMN stages on day 28 and day 50 (Supplementary Fig. 5A). However, the proportions of different AS event types did not differ significantly across developmental stages or ALS subgroups (Fig. 7B).

**Figure 7.**
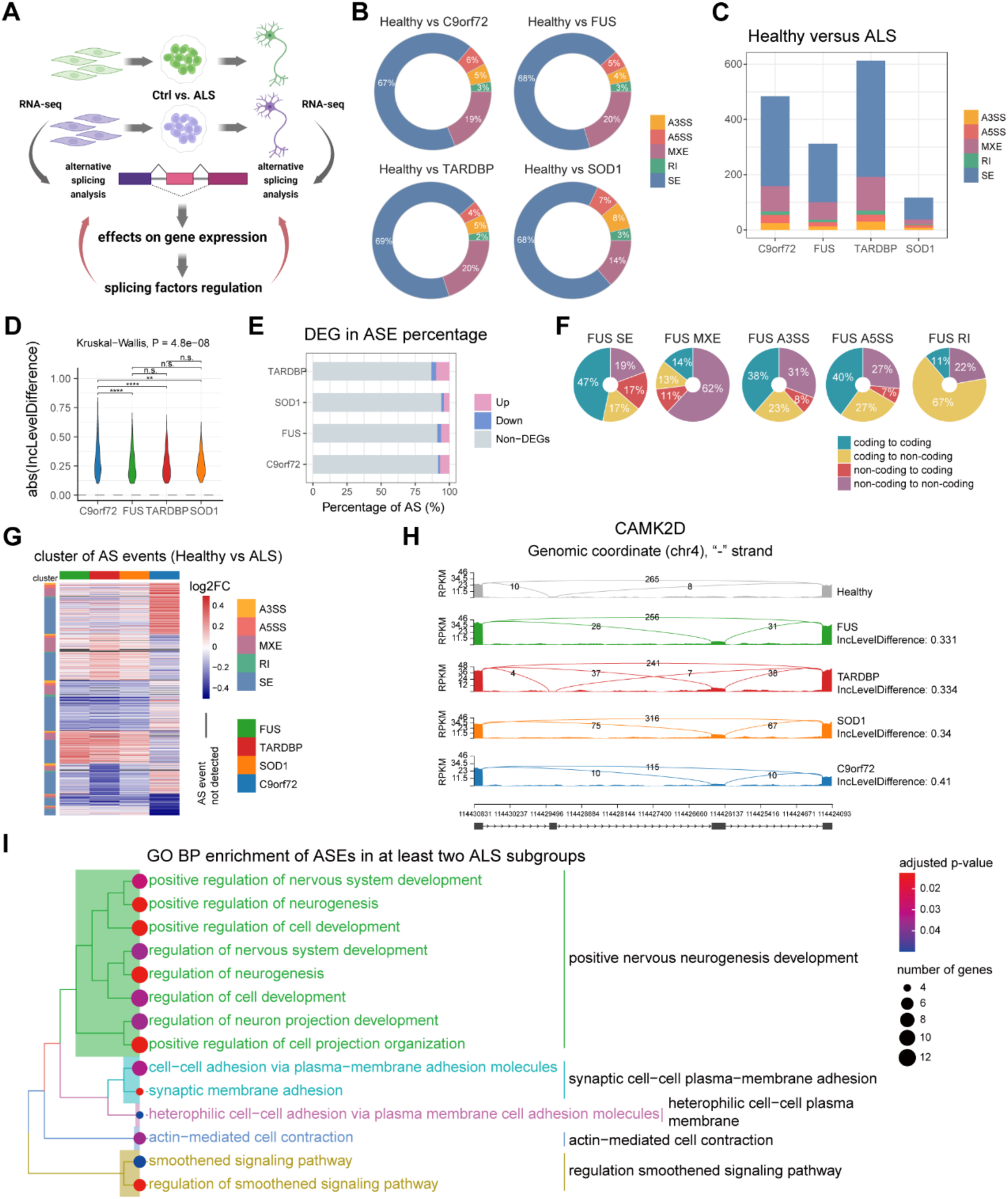
Alternative splicing alterations in ALS patients’ D50-iMNs. (A) Schematic illustrating the analysis workflow of alternative splicing. (B) Proportions of splicing types in ALS D50-iMNs compared to healthy D50-iMNs (FDR < 0.05, |InclusionLevelDifference| > 0.1; Wilcoxon signed-rank test; * p < 0.05, ** p < 0.01, *** p < 0.001, **** p < 0.0001, n.s., not significant). (C) Numbers of splicing types in four different ALS subgroups compared to healthy samples. (D) Distribution of absolute inclusion level differences in the four ALS subgroups (Wilcoxon signed-rank test; * p < 0.05, ** p < 0.01, *** p < 0.001, **** p < 0.0001, n.s., not significant). (E) The ratio of up- and down-regulated genes and non-DEGs by ALS D50-iMNs. (F) The distribution of annotated transcripts changing on potential protein-coding ability influenced by AS events in ALS-*FUS* D50-iMNs. Coding potentials are calculated by mapping splicing events to transcripts. Chart areas are proportional to changing numbers of annotated transcripts for each AS type. (G) The heatmap shows inclusion level differences of AS events significant in at least one of the four ALS subgroups from included (red) to excluded (blue) (1308 events in 929 genes). AS events not detected as significant are depicted as dark grey. (H) The sashimi plot of *CADM1* and *CAMK2D* shows significant AS events in ALS subgroups compared to healthy lines. Lower box depicting gene structure. Arrows indicate the direction of gene transcription. (I) Tree plot depicting enriched BP terms of genes showing AS significant changes in at least two ALS subgroups.

Interestingly, we observed the most significant changes in AS events in iMNs carrying *TARDBP* mutations, followed by *C9orf72* and *FUS* mutations on day 50 (Fig. 7C; Supplementary Fig. 5D). We identified 312, 613, 117, and 484 AS events in ALS-*FUS*, ALS-*TARDBP*, ALS-*SOD1* and ALS-*C9orf72* on day 50, respectively, and the distribution of alternative splicing changes (exon inclusion levels) was similar among all ALS subgroups (Supplementary Fig. 5A, 5B). On day 50, about 10% of the genes showed differential expression between ALS subgroups and healthy controls (Fig. 7E), with the proportion of up-regulated genes being higher than that of down-regulated genes (Fig. 7E).

To better understand the impact of alternative splicing on protein function, we investigated the coding potential of the transcribed sequence. We mapped transcripts to splicing events to explore possible protein features affected by splicing. Our analysis showed that changes in protein-coding ability were mainly preserved in SE and MXE events (Fig. 7F; Supplementary Fig. 5C). However, RI events caused significant changes in protein-coding, converting them from non-coding to coding or vice versa (Fig. 7F; Supplementary Fig. 5C).

Next, we analyzed shared AS changes across different ALS subgroups. To determine the significant inclusion level differences of AS events, we calculated Pearson’s correlation coefficients and found a strong correlation among ALS-*FUS*, ALS-*TARDBP* and ALS-*SOD1* (Supplementary Fig. 5E). We then grouped significantly differentially spliced events into seven distinct clusters (Fig. 7G) and identified 1308 AS events in at least one ALS subgroup on day 50 (Fig. 7G). Of these, three were significantly changed in all four ALS subgroups, 28 in three ALS subgroups, and 153 in two ALS subgroups.

Our study identified *CAMK2D* and *CADM1* as significant players in all ALS subgroups (Fig. 7H; Supplementary Fig. 6A, 6B). CAMK2D, a member of the serine/threonine protein kinase family and the Ca^2+^/calmodulin-dependent protein kinase subfamily, is known for its crucial role in various aspects of plasticity at glutamatergic synapses^51^. On the other hand, cell adhesion molecule 1 (CADM1) is a crucial facilitator of cell adhesion, highly expressed in the human prefrontal lobe and implicated in the genetic architecture of ADHD^52,53^. In mature ALS iMNs, *CADM1* had SE events, and *CAMK2D* had MXE events across all ALS subgroups.

We then focused on identifying changes in AS events in at least two ALS subgroups. Our analysis showed that the genes with AS changes are mainly localized in the presynaptic area and axon (Supplementary Fig. 5F), as well as enriched in the regulation of nervous system development, such as synaptic membrane adhesion and regulation of nervous system development (Fig. 7I). These findings suggest that mutations in ALS-causative genes affect the splicing events of genes related to essential neuronal functions, including synapse regulation in iMNs on day 50.

### Splicing factors dysregulates AS events on transcription level in mature iMNs

Transcription factors (TFs) are crucial in controlling gene expression at the transcription level. Recent studies have shown that most splicing occurs during transcription, which has led to the discovery that the chromatin state affects alternative splicing^54^. Therefore, it is plausible that transcription factors can influence splicing outcomes^54^. To investigate the role of TFs in splicing regulation in ALS iMNs on day 50, we used ChIP-X Enrichment Analysis Version 3 (ChEA3) to identify the TFs with aberrant splicing events^55^. KMT2A was found to regulate ASEs in *FUS*, *TARDBP*, and *SOD1* iMNs (Supplementary Fig. 7A, B). KMT2A is a transcriptional coactivator that is essential in regulating gene expression. Recent evidence suggests that MeHg, a toxic compound, accelerates necroptotic cell death in SOD1-G93A cells by increasing REST gene expression via Sp1/KMT2A complex^56^.

RNA-binding proteins (RBPs) also play an important role in regulating gene expression by binding to short RNA sequences. They are crucial in several RNA-related processes, such as pre-mRNA splicing, cleavage and polyadenylation, RNA stability, RNA localization, RNA editing, and translation^57–59^. Dysfunctions in RBP are associated with various genetic and somatic disorders, including neurodegeneration^60^. Recent studies highlight the essential role of RBPs in RNA metabolism and their involvement in ALS pathogenesis^61^.

To understand the molecular mechanisms underlying abnormal AS in ALS subgroups mediated by RBPs, we conducted a binding motif enrichment analysis on 91 known RNA binding proteins using the rMAPS motif tool. We evaluated multiple regions for each AS event type. Motif analysis showed that 60 RBPs bound with accompanying AS events in at least one ALS subgroup on day 50 (Fig. 8B).

**Figure 8.**
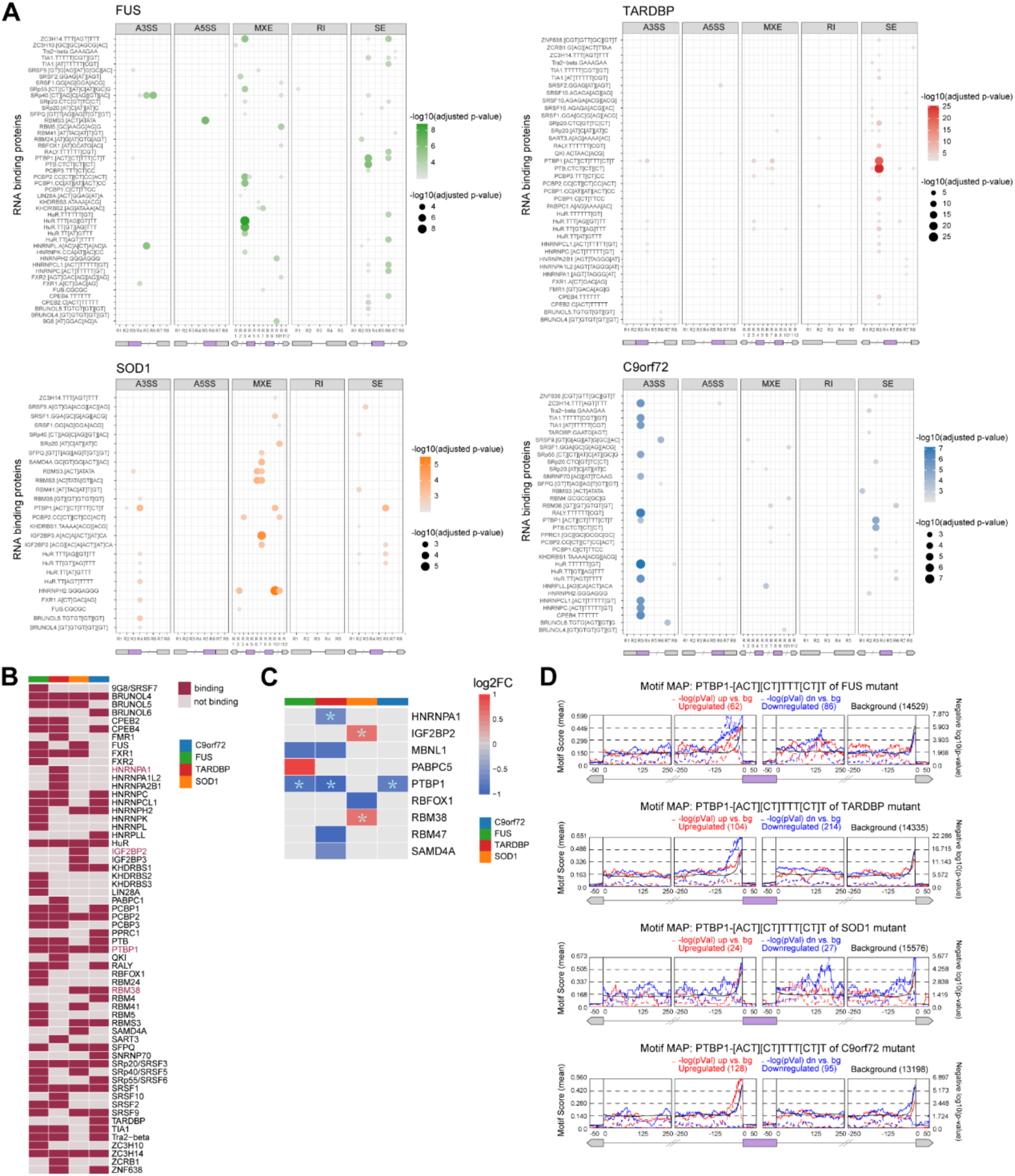
Splicing factors regulation of aberrant AS events in ALS iMNs on day 50. (A) Bubble plot of significantly enriched RBPs’ binding motifs around AS events on day 50 (adjusted p-value < 0.001). The region (R) of each splicing event is numbered from R1 to R(n), 5’ to 3’, where n is the total region for each event type (bottom). Purple boxes represent exons of interest for each event type; gray boxes represent up- and downstream exons. Lines separated with slashes represent introns. Intronic regions consist of 250-bp flanking sequences, and exon regions consist of 50-bp sequences from the start or end of the exon, known for frequent splicing factor binding. (B) The heatmap for motif binding overlaps of RBPs across all ALS subgroups on day 50. (C) The heatmap showing log2 fold change value of significant differentially expressed RBPs used in rMAPS between ALS and healthy iMNs on day 50. RBPs marked by “*” are DEGs with binding sites in regions flanking alternative splicing events of the corresponding ALS subgroup. (D) Positional distribution of the PTBP1 binding motif. The PTBP1 binding motif is shown above the panel. The purple box represents the skipped exon, the left gray box represents the 3’ end, and the right gray box represents the 5’ end of the exon. Lines represent 250-bp intronic sequences before and after each shown exon. The solid lines show the mean motif score calculated as density within a 50-bp sliding window, representing the percentage of nucleotides covered by the PTBP1 binding motif. The black line indicates the background signal in this region. Red and blue lines depict the enrichment of PTBP1 binding motifs around exons more or less utilized by ALS subgroups. The dash lines show the −log10(p-value) of the motif enrichment in these windows, with higher peaks indicating a more significant p-value for that region. Significance was determined by comparison to a ‘background set’ of exons without splicing changes in expressed genes (bottom).

We have observed that RNA binding motifs (RBM) are mainly enriched in various AS events specific to ALS subgroups. For ALS-*FUS* and ALS-*SOD1*, RBPs’ binding motifs were found around intron regions in MXE and SE events. In the case of ALS-*C9orf72*, the R3 region of A3SS events exhibited a higher enrichment of RBMs compared to other AS event types (Fig. 8A). Out of the 60 identified RBPs, five of them are reported as genes or genetic variants associated with ALS, according to ALSoD. These include FUS, HNRNPA1, HNRNPA2BA, TARDBP, and TIA1^62^ (Fig. 8B). Specifically, we identified BRUNOL4, HuR, PCBP2, PTBP1, SRSF1, SRSF3, and ZC3H14 as having binding site motifs flanking an AS event, or more times across all ALS subgroups (Fig. 8B).

We made a significant discovery: HNRNPA1 binding sites were specific to ALS-*TARDBP*, while IGF2BP2 and RBM38 were specific to ALS-*SOD1*. *PTBP1* exhibited approximately 1-fold down-regulation in ALS-*FUS*, ALS-*TARDBP*, and ALS-*C9orf72*. The binding motif of PTBP1 was enriched in regions flanking significant AS events across all ALS subgroups (Fig. 8C, D). Our RBM analysis revealed that ES events in iMNs were typically enriched for binding sites of PTBP1 within 100 nucleotides upstream of the skipped exon or 200 nucleotides downstream of the skipped exon (Fig. 8A, D). PTBP1, a splice factor belonging to the ubiquitously expressed heterogeneous nuclear ribonucleoproteins (hnRNPs) subfamily, exhibited significant downregulation^63^. This finding highlights the potential role of PTBP1 downregulation in neurogenesis and neurodegenerative disease^64^, necessitating further research.

Recent studies have also shown that mutations in hnRNPs have been linked to neurodegenerative diseases, especially ALS and FTD^65^. We also analyzed RBPs reported in previous studies to investigate how their expression changed on day 50^57^. Most of these RBPs exhibited significant down-regulation on day 50, suggesting dysfunction in RBPs and their potential contribution to dysregulated AS in ALS iMNs. Our findings suggest that differentially expressed RBPs or other upstream transcription regulators in ALS iMNs may affect abnormal AS events.

## DISCUSSION

Researchers have conducted various transcriptome and proteomics studies on iMNs derived from ALS patients to identify pathological features of the disease. However, most of these studies only looked at transcriptome changes in mature iMNs, which could not explain the molecular mechanisms of progressive degeneration of motor neurons. Previous studies have focused on one or two ALS genes, leaving us with a limited understanding of the different ALS cases’ divergent and convergent pathological mechanisms. In this study, we have systematically characterized transcriptomic changes in ALS iMNs with *C9orf72*, *FUS*, *TARDBP*, and *SOD1* mutations at 4 different temporal time points (Fig. 1). This provides us with experimental insight into cellular pathways underlying the progression of ALS and offers insights into the pathomechanisms by which motor neurons progressively degenerate.

By mapping representative GO terms enriched from NPCs to mature iMNs, we have elucidated the representative biological processes in mature iMNs that are disrupted by each ALS gene (Fig. 3). These findings not only indicate the specific pathological mechanisms involved in different ALS genes, but also suggest that the treatment of ALS should be tailored to the genetic cause of ALS patients, offering a promising avenue for personalized medicine.

We discovered several biological process categories that were significantly enriched in more than three ALS subgroups in mature iMNs, including actin filament-based process, cell-matrix adhesion, signal transduction by p53 class mediator, synapse organization and axonogenesis (Fig. 3E). Additionally, these data indicates that dysregulation of cytoskeleton, cell adhesion, cellular composition organization, synaptic function, cellular response, and neuronal development may begin early in ALS and persists in later stages of the disease, playing essential roles in the complex pathology networks (Fig. 3E).

Additionally, we used PROGENy to examine the signaling pathways involved in motor neuron development and maturation. Our analysis revealed the activation of inflammation-related NF-κB and TNF-α pathways in ALS iMNs on day 50 (ALS-*FUS* and ALS-*SOD1*). Notably, the activation was less pronounced in ALS-*SOD1* compared to ALS-*FUS* (Fig. 5F). The potential role of neuroinflammation has gained significant attention in ALS research. Our findings suggest that ALS genes may promote neuroinflammation at the transcriptome level in mature iMNs.

Previous research has reported dynamic changes in AS in mammalian brains and human iPSC-based models during developmental stages or in mature motor neurons^39,66,67^. However, our understanding of the regulatory mechanisms of AS across motor neuron differentiation shared by different mutants is currently limited. This study investigated AS events and examined the different regulatory mechanisms during iMN development and maturation. From day 28 to day 50 of iMN maturation, genes with abnormal AS events were functionally over-represented in neuron projection development and membrane depolarization during action potential. These genes were also significantly present in cellular components related to the synaptic membrane, axon initial segment, and ion channel complex, which are central to ALS motor neuron pathophysiology (Fig. 6D). Furthermore, genes exhibiting AS only in healthy subgroups, but not in any ALS subgroups, were associated with pre-synapse or post-synapse identified in SynGO^49^ (Fig. 6E, F). One specific example is the Neurexins (NRXN1, NRXN2 and NRXN3), a family of proteins that serve as cell adhesion molecules and receptors in the vertebrate nervous system. Mutations in the neurexins family have been identified in patients with autism spectrum disorder (ASD) and schizophrenia (SCZ)^68^.

Specifically, we found that ALS genes induce more aberrant AS events in mature iMNs, a finding that could have profound implications for our understanding of the disease. Furthermore, it is worth noting that most of these events did not necessarily affect the gene expression at a significant level (Fig. 7E, F), suggesting that assessing only gene expression without considering actual isoform usage can provide limited information about the transcriptomic changes that affect disease biology.

Our analysis identified MXE events in *CAMK2D* on day 50 (Fig. 7H; Supplementary Fig. 6B). In mammalian cells, the CAMK2 enzyme consists of four different chains: alpha, beta, gamma, and delta. The *CAMK2D* gene produces the delta chain of the enzyme. CAMK2 is critical in calcium signaling, synaptic plasticity and memory formation^51^. CAMK2A or CAMK2B are the most abundant CAMK2 protein family isoforms in the brain, and the role of these two isoforms in neuronal functioning, such as learning and plasticity, has been studied in animal model^51^. Recently, the importance of CAMK2A and CAMK2B for normal human neurodevelopment was also demonstrated^69–72^. A significant alteration in splicing patterns of *CAMK2D* in all ALS iMNs may represent an innovative target for further studies. Additionally, our data revealed differences in the exon 8 usage of *CADM1* isoforms between ALS subgroups and controls (Supplementary Fig. 6A). We observed reduced exon 8 inclusion in ALS-*FUS*, ALS-*TARDBP*, and ALS-*SOD1* subgroups, while ALS-*C9orf72* showed elevated exon 8 inclusion (Supplementary Fig. 6A).

Recent discoveries using in vivo and in vitro models of ALS have revealed that early synaptic dysfunction occurs at the onset of the disease, prior to motor neuron degeneration symptom manifestation^73^. This observation is further supported by post-mortem analysis of an ALS patient’s tissues, where no degeneration of lower motor neurons (LMNs) was detected in their corticospinal tract and minimal axon degeneration in the spinal cord^74^. Further studies have shown that, by examining the electrophysiological properties of LMNs, ALS patients in the early stages of the disease exhibit signs of corticospinal degeneration, loss of LMNs, and altered excitability of surviving motor units, while NMJs remain functional^43,73,75–79^.

Based on these findings, we noticed that genes with AS changes involved in at least two ALS subgroups on day 50, demonstrate significant enrichment in synaptic dysfunction (Fig. 7I; Supplementary Fig. 5F). Furthermore, the temporal splicing alterations caused by ALS genes align with the electrophysiological properties illustrated in Fig. 2, and the impacted molecular pathways during iMN maturation depicted in Fig. 6. This indicates potential deficits in synaptic function at different stages of iMN maturation, including the regulation of alternative splicing and gene expression.

Our analysis of motif enrichment around AS regions has identified several RBPs as potential factors that bind to them (Fig. 8B). Mutations in hnRNPs have previously been linked to neurodegenerative diseases, particularly ALS and FTD^80^. In neurodegeneration, TDP-43 and FUS are the most well-known hnRNPs^80^. Specifically, the IPA analysis revealed that some hnRNPs have direct, experimentally confirmed interactions with the key pathological genes and proteins associated with ALS, including TDP-43, C9orf72, FUS and Tau^80^. It is important to note that all ALS data sets displayed differences in the expression of several previously identified RBPs (Fig. 8C). One such RBP is PTBP1, which is enriched in regions flanking significant AS events and was dramatically down-regulated in ALS-*FUS*, ALS-*TARDBP*, and ALS-*C9orf72* (Fig. 8C). PTBP1 is an RNA-binding protein and splicing regulator that is broadly expressed in non-neuronal and neuronal progenitor cells and represses neuronal-specific alternative splicing^81–83^. Although PTBP1 belongs to hnRNPs as a splicing regulator and regulates the activity of other transcription factors during neurogenesis and neuronal differentiation, it is still unknown whether PTBP1 may or may not cause motor neuron degeneration in ALS and requires further evaluation^28,80,84–86^. A previous study showed that PTBP1 was complexed with FUS and identified as one of the FUS interactors, explaining its inability to inhibit pre-mRNA splicing in FUS-immunodepleted extract^87^. Hence, PTBP1 may contribute to neuronal death or perturb other mechanisms that maintain healthy neuronal function through interaction with other ALS-associated proteins.

In summary, our research has identified changes in the transcriptome of ALS during motor neuron development and disease progression and has created a comprehensive map of alternative splicing. These findings contribute to the growing evidence implicating dysregulated synaptic functions occurring before motor neuron degeneration in ALS and being a shared molecular characteristic across different ALS genes. They will also serve as a valuable resource for understanding ALS pathogenesis’s complexity, heterogeneity, and diversity.

## Methods

Methods and any associated references are available in the online version of the paper. Note: Supplementary Information includes 8 figures, the list of the primers and antibodies used in this study is available online.

## Consent for publication

Not applicable.

## Availability of data and materials

The datasets during and/or analysed during the current study available from the corresponding author on reasonable request.

## Competing interests

The authors declare no competing interests.

## Acknowledgements

We thank all members of the Wang lab for helpful discussions.

## Funding

This work is supported by the National Key R&D Program of China (Grant No2018YFA0107903), and the Shanghai Municipal Science and Technology Major Project (Grant No.2019SHZDZX02), Shanghai Key Laboratory of Aging Studies [19DZ2260400 to W. W], Guizhou Provincial Higher Education Science and Technology Innovation Team ([2023]072)

## Author’s Contribution

G.M.M, C.C.X and B.Y.L designed and performed the experiments and analyzed data; L.J helped with cell culture; all other authors provided technical support; W.Y.W, G.Q.Y, Y.W.M, and Y.C supervised the project.

## Materials and methods

### Generation and culture of iPSCs

Fibroblasts from ALS patients carrying ALS mutations and healthy controls were obtained from the Coriell Institute for Medical Research and ATCC. Details of the lines used in this study are provided in Table S1. These fibroblasts were reprogrammed into iPSCs with the non-integrating Sendai virus (Life Technologies). The reprogrammed iPSCs were maintained on matrigel (Corning) with Essential 8 Medium media (Life Technologies), and then were passaged using EDTA (Life Technologies, 0.5 mM). The iPSCs were cultured at 37 °C at 5% CO_2_ and 5% O_2_.

### Motor neuron differentiation

The motor neuron differentiation protocol was adapted from a previously published protocol^1^. In brief, iPSCs were split 1:4 on Matrigel-coated plates, and then differentiated to NPCs in chemically defined N2B27 medium, consisting of DMEM/F12, Neurobasal medium at 1:1, 0.5× N2, 0.5× B27, 0.1mM ascorbic acid (Sigma), 1× Glutamax and 1× penicillin/streptomycin, β-mercaptoethanol (all from Life Technologies). 0.2 µM LDN-193189 (Selleck Chemicals), 2 µM SB431542 (Selleck Chemicals), and 3 µM CHIR99021 (Selleck Chemicals) were added in the medium for 6 days for NPC induction. On day 7, NPCs were dissociated with Dispase (1 mg/mL) and split at 1:2. Then NPCs were cultured under condition with 1 µM retinoic acid (RA, Sigma), 1 µM Smoothened Agonist (SAG, Selleck Chemicals), 1 µM CHIR99021, 0.2 µM LDN193189 and 2 µM SB431542 in N2B27 medium for another 6 days and differentiated into OLIG2+ motor neuron precursors (MNPs). OLIG2+ MNPs were cultured in suspension in the N2B27 medium with 1 µM RA and 1 µM SAG for 6 days and differentiated into ISL1+ MNs. The ISL1+ MNs were dissociated into single cells with 0.25% trypsin (Life Technologies) and then plated on poly-L-ornithine (10 mg/mL, Sigma)/laminine (5 mg/mL, Life Technologies) coated plates. After 10 days cultured with 1 µM RA, 0.1 µM Pur and 0.1 µM Compound E (Millipore), 10 µg/mL BDNF, 10 µg/mL GDNF and 10 µg/mL CNTF MNs were differentiated into mature into CHAT+ MNs. Then, MNs were cultured in N2B27 for 22 days without any addition and then collected for experiments.

### Neuromuscular junction innervation

The neuromuscular junction detection protocol was adapted from a previously published protocol^1^. The glass coverslips were treated with trimethoxysilylpropyldiethylenetri-amine (DETA, Sigma), followed the published protocol^2^. C2C12 cells were culture on the treated glass coverslips with Matrigel-coated. C2C12 cells were cultured in DMEM with 10% FBS, and then were induced to form myotube by switching to DMEM containing 10% horse serum. Day 18 MNs were digested into single cells and plated on the induced myotubes for 10 days, after which the neuromuscular junctions were visualized by performing immunofluorescence of CHAT (1:50, Millipore) and α-BTX-594 (1:200, Sigma) staining.

### Immunofluorescence

Cells cultured on glass coverslips were fixed with 4% paraformaldehyde in 1× phosphate buffer saline (PBS) for 10 min at room temperature, and then blocked with 10% normal donkey serum (Life Technologies) with 0.2% TritonX-100 in PBS for one hour. Then cells were then incubated with primary antibodies in 1% BSA + 0.1% TritonX-100 in PBS overnight at 4 °C and incubated with secondary antibodies for one hour at room temperature. The following primary antibodies were used: Rabbit anti-OCT-4 (1:400; Cell Signaling; #2840), Rabbit anti-SSEA-4 (1:500; Cell Signaling; #4755), Rabbit anti-NANOG (1:250; Abcam; ab109250), Rabbit anti-PAX6 (1:1000; Abcam; ab154253), Mouse anti-NESTIN (1:500; Abcam; ab18102), Rabbit anti-OLIG2 (1:1,000; Millipore; AB9610), Rabbit anti-ISLET1 (1:1,000; Abcam; ab109517), Mouse anti-SMI31 (1:1,000; Covance; SMI-31P), Mouse anti-SMI32 (1:1,000; Covance; SMI-32P), Mouse anti-HB9 (1:1,00; DSHB; 81.5C10), Rabbit anti-TUJ1 (1:1,000; Abcam; ab14545), Goat anti-CHAT (1:200; Millipore; AB144P) and α-BTX-594 (1:200; Sigma; T0195-.5MG).

### Electrophysiology

Whole-cell patch-clamp recordings were applied to detect the firing properties of iPSC-derived MNs on day 50. The protocol of whole-cell patch-clamp recordings was adapted from previously published work^3^. The artificial cerebral spinal fluid (aCSF) we used consisted of: 119 mM NaCl, 5 mM KCl, 1.25 mM NaH_2_PO4.2H_2_O, 26 mM NaHCO_3_, 2 mM CaCl_2_, 1 mM MgSO_4_, 5 mM glucose and 95% O_2_/5% CO_2_.

Recording pipettes were filled with K-gluconate-based current clamp internal solution containing 130 mM K-gluconate, 10 mM KCl, 10 mM HEPES, 0.2 mM EGTA, 0.5 mM Na_3_-GTP, 4 mM Mg_2_-ATP, 10 mM Na-phosphocreatine, pH 7.2 and 290 mOsm. Electrophysiological data were analyzed using Clampfit10 software (Axon Instruments). Differences between healthy controls and ALS subgroups were analyzed with student’s t-test. P < 0.05 was considered significant. Data of Na^+^ and K^+^ currents were presented as mean ± SEM.

### RNA extraction and qPCR

Total RNA from iPSCs, NPCs, D28-iMNs and D50-iMNs was extracted using TRIzol (Invitrogen). DNA digestion and reverse transcription were performed using the Hifair™ **Ⅲ** 1st Strand cDNA Synthesis SuperMix for qPCR (YEASEN) following the manufacturer’s instructions. qPCR was performed on cDNA using qPCR SYBR Green Master Mix (UNIQ) with the QuantStudio 7 Flex Real-Time PCR System according to manufacturer’s instructions. The relative expression levels of target genes were measured using the ddCt method normalizing to control genes expression. ACTIN was used as reference in ALS-*SOD1*, ALS-*FUS*, and ALS-*TARDBP* subgroups. TFRC was used as reference in ALS-*C9orf72*. Primers used in qPCR are listed in Table S2.

### RNA sequencing and differential gene expression analysis

Total RNA was extracted from iPSCs, NPCs, D28-iMNs and D50-iMNs using TRIzol (Invitrogen). RNA purification, reverse transcription and library construction were performed in Mingma Technologies Co., Ltd at Shanghai according to the manufacturer’s instructions. RNA concentration and integrity were measured using Nanodrop (Thermo Scientific) and an Agilent 2100 Bioanalyzer (Agilent). The mRNA-focused sequencing libraries from total RNA were prepared using VAHTS mRNA-seq v3 Library Prep Kit(VAHTS, NR611). PolyA mRNA was purified from total RNA and then fragmented. The products were enriched with PCR amplification to create the final cDNA libraries. After library construction, Qubit 3.0 fluorometer dsDNA HS Assay (Thermo Fisher Scientific) was used to quantify concentration of the resulting sequencing libraries, while the size distribution was analyzed using Agilent BioAnalyzer (Agilent). The libraries were sequenced with Illumina HiSeq PE150 sequencing system following Illumina-provided protocols in Mingma Technologies Co., Ltd at Shanghai. After removing the adapter sequences and low-quality reads using Trimmomatic (v.0.39)^4^ and quality checking the data using FastQC (v.0.11.9; http://bioinformatics.babraham.ac.uk/projects/fastqc/), the reads were aligned to the GRCh37 reference genome (build GRCh37.p13/hg19) with HISAT2 (v.2.2.1). The gene-wise read-counts were quantified from the aligned reads by FeatureCounts in Rsubread (v.2.0.1)^5^ using GENCODE GTF annotation version 19. For hierarchical clustering, the normalized data were calculated based on Manhattan correlations with Ward’s method. Differential expression analysis of normalized gene expression was performed using DESeq2 (v.1.34.0)^6^, with fold change ≥ 1.5 and false discovery rate (FDR) < 0.05 indicating significant DEGs. Transcriptome alteration pattern across iMN developmental stages of DEGs identified in day 50 was clustered by utilizing the function cutreeDynamic from the R package dynamicTreeCut (v.1.63-1)^7^. PROGENy signalling pathway activities were estimated using the decoupleR (v.2.0.1)^8^ and progeny (v.1.16.0)^9^ packages in R.

### Alternative splicing analysis

RNA-seq data were then used to perform differential splicing analysis using rMATS (v.4.1.2)^10^. We used the reads on target and junction counts (JCEC) for each AS event. The significant AS events were filtered and categorized based on a IncLevelDifference (ΔPSI) absolute value greater than 0.1 and FDR < 0.05. The splicing event categories included SEs, A5SSs, A3SSs, MXEs, and RIs. Sashimi plots of AS events were generated using rmats2sashimiplot (v.2.0.3). Binding motif enrichment analysis was performed by rMAPS2 (v2.0.0)^11^ using rMATS output. Known binding motifs for RNA-binding proteins were covered in rMAPS2. Significantly spliced regions were used as the target regions for motif enrichment, and not significantly spliced regions were used for estimating background binding levels. The lengths of the intronic and exonic regions were assessed and plotted using 250 bp and 50 bp, respectively. Changes in the coding potential of the target genes which were alternative spliced in ALS subgroups were estimated using R package Mapping Alternative Splicing Events to pRoteins (MASER, v.1.12.1; https://github.com/DiogoVeiga/maser). MapTranscriptsToEvents() function in MASER was used to identify transcripts potentially affected by alternative splicing events. All transcripts attached to alternatively spliced regions were annotated using GENCODE GTF annotation version 19.

### Enrichment analysis

Gene ontology (GO) analysis was performed using clusterProfiler R package (v.4.2.1)^12^, and adjusted p-value < 0.05 was considered as significant. Treeplot() function in clusterProfiler performed hierarchical clustering of GO enriched terms. The overrepresentation of synaptic GO terms was estimated by the SynGO online portal (www.syngoportal.org)^13^.

### Gene-transcription factor interaction analysis

The ChEA3 web browser^14^ were used to infer upstream regulators of genes with differentially alternative spliced. Mean rank was used in this study.

### Statistical analyses

The statistical analyses used can be found in the figure legends. All statistical tests were computed using GraphPad Prism or R Studio. GraphPad Prism 9 was used to visualize data and calculate statistics for grouped analyses. Data are depicted as mean ± standard error of the mean (SEM) in the case of n ≥ 3, as noted in the figure legends. For correlation analyses Pearson correlations were employed. The Kruskal-Wallis test with Dunn’s correction for the ALS variables was performed for the distribution of exon inclusion levels. Function pairwise.wilcox.test() was used to calculate pairwise comparisons between group levels with corrections for multiple testing. Schematic illustrations were created with Biorender (https://biorender.com/). Data visualization of sequencing data analyses was done in R using the VennDiagram (v.1.7.3), UpSetR (v.1.4.0), pheatmap (v.1.0.12), circlize (v.0.4.15), corrplot (v.0.92), lessR (v.4.3.0), scatterpie (v.0.2.1), and ggplot2 (v.3.4.2) packages available from CRAN.

## Supplementary information

**Supplementary Fig 1.**
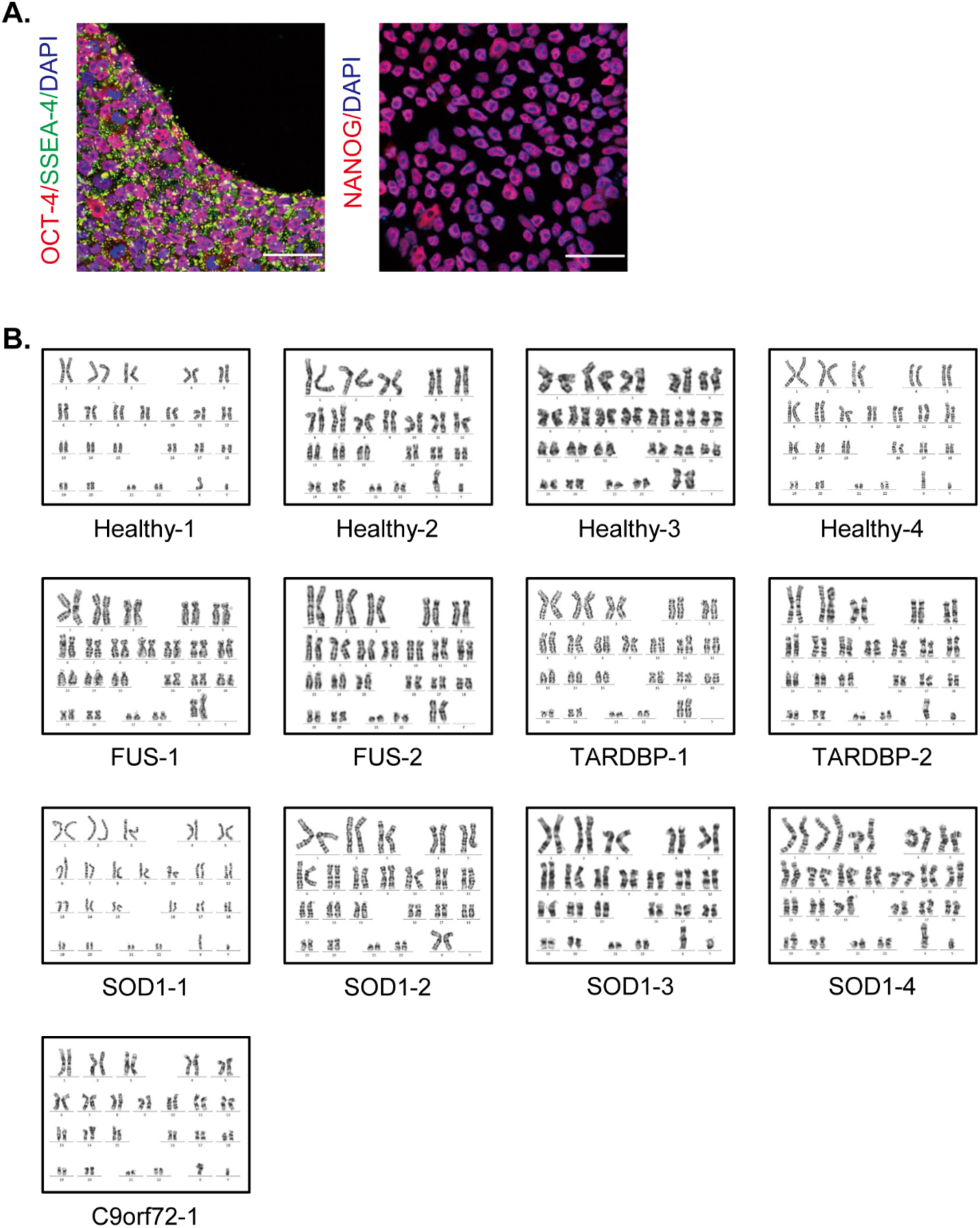
Characterization of iPSCs. (A.) Immunofluorescence of pluripotent markers (OCT4, SSEA-1, and NANOG) for iPSCs. The scale bar represents 50 μm. (B.) G-band karyotype analysis of iPSCs from controls and ALS patients.

**Supplementary Fig 2.**
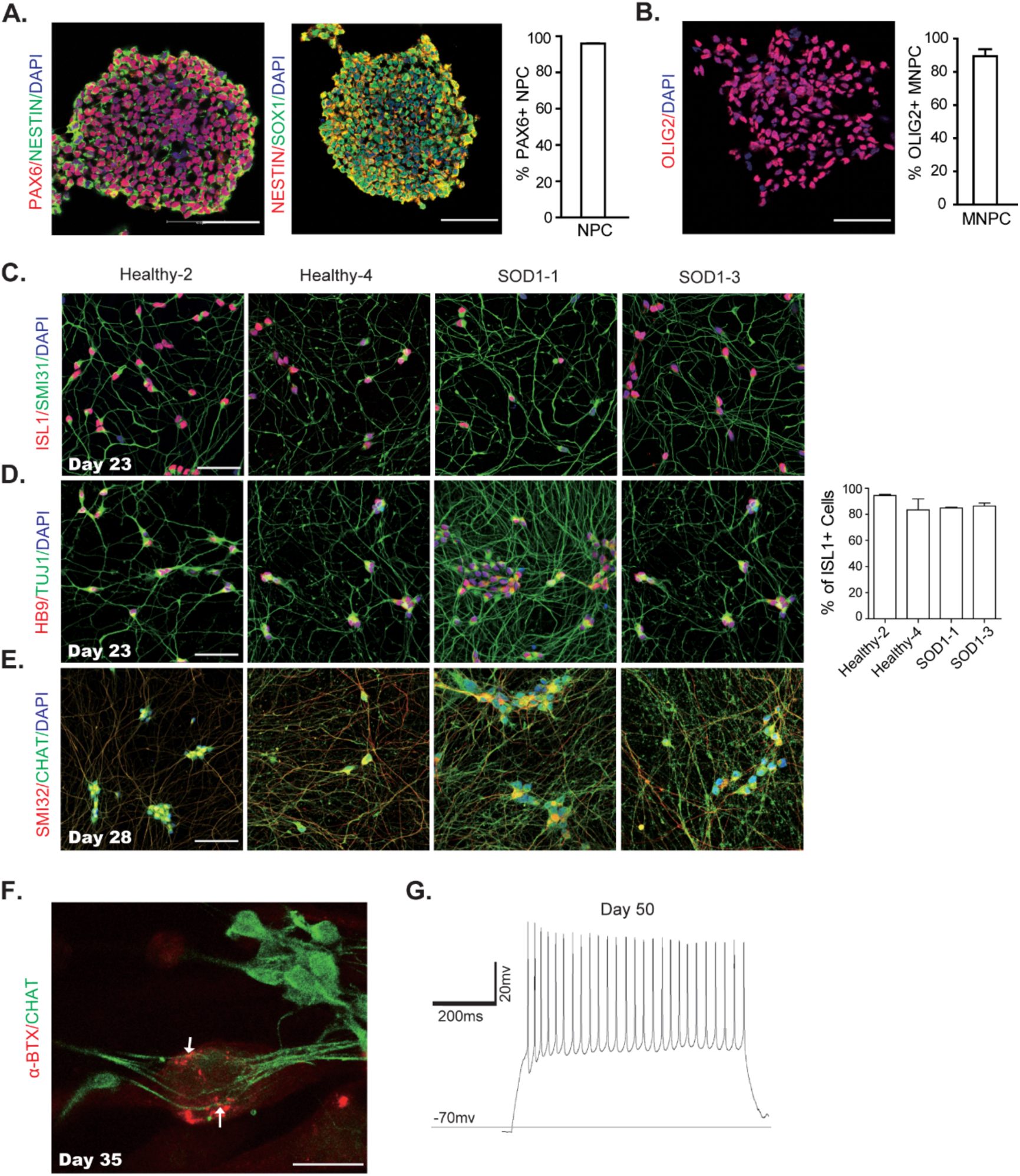
Characterization of iMN differentiation. (A.) Representative images of SOX1+/PAX6+/NESTIN+ NPCs after 6 days of culture in CHIR + SB + LDN condition. Cell nuclei were stained with DAPI (blue). Quantification of PAX6+ cells was shown. The scale bar represents 50 μm. (B.) Representative images of pure OLIG2-positive motor neuron progenitors (MNPs) on day 12. Cell nuclei were stained with DAPI. The scale bar represents 50 μm. (C-E.) Representative images of the two motor neuron transcription factors ISL1/HB9, the functional motor neuron marker CHAT/SMI32 and the pan neuronal marker TUJ1. The scale bar represents 50 μm (left). The average percentage of ISL1+ motor neurons in healthy controls and ALS lines. No statistically significant difference was found (mean ± SEM; n = 3 independent experiments; student’s t-test; n.s., not significant) (right). (F.) Representative images of iMNs, stained with the CHAT antibody (green), formed neuromuscular junctions labeled with α-bungarotoxin (α-BTX, red) when co-cultured with myotubes. The scale bar represents 20 μm. (G.) An Example voltage trace illustrates the response to current injections and highlights the generation of APs triggered by depolarizing current injections.

**Supplementary Fig 3.**
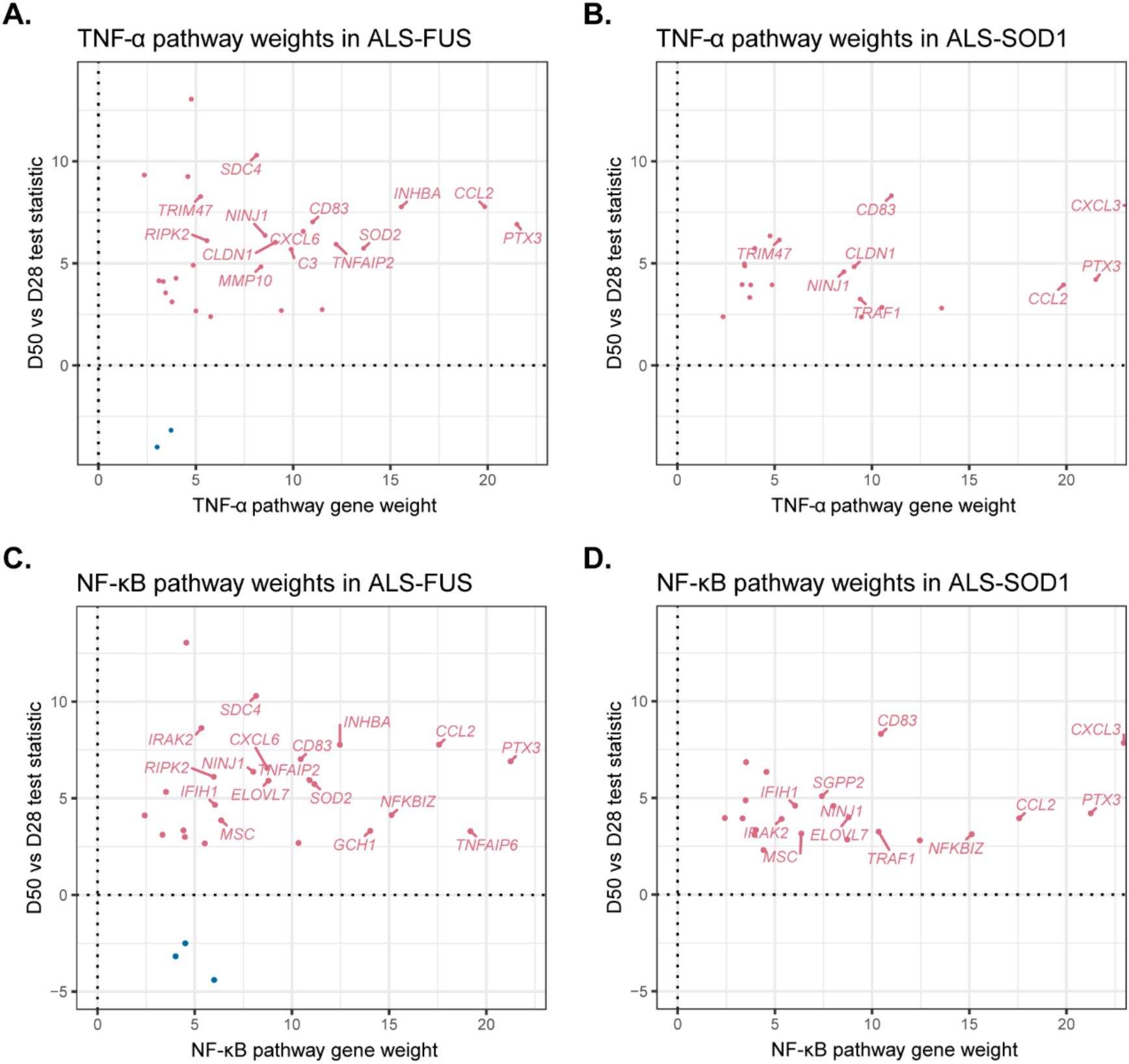
Expression changes of selected pathway genes in ALS D50-iMNs versus D28-iMNs. (A-D.) Expression changes of TNF-α pathway genes in ALS-*FUS* (A), ALS-*SOD1* (B), and NF-κB pathway genes in ALS-*FUS* (C), ALS-*SOD1* (D) in D50-iMNs versus D28-iMNs according to their PROGENy weights. Genes in D50-iMNs increasing pathway activity are colored red, and genes decreasing pathway activity are colored blue.

**Supplementary Fig 4.**
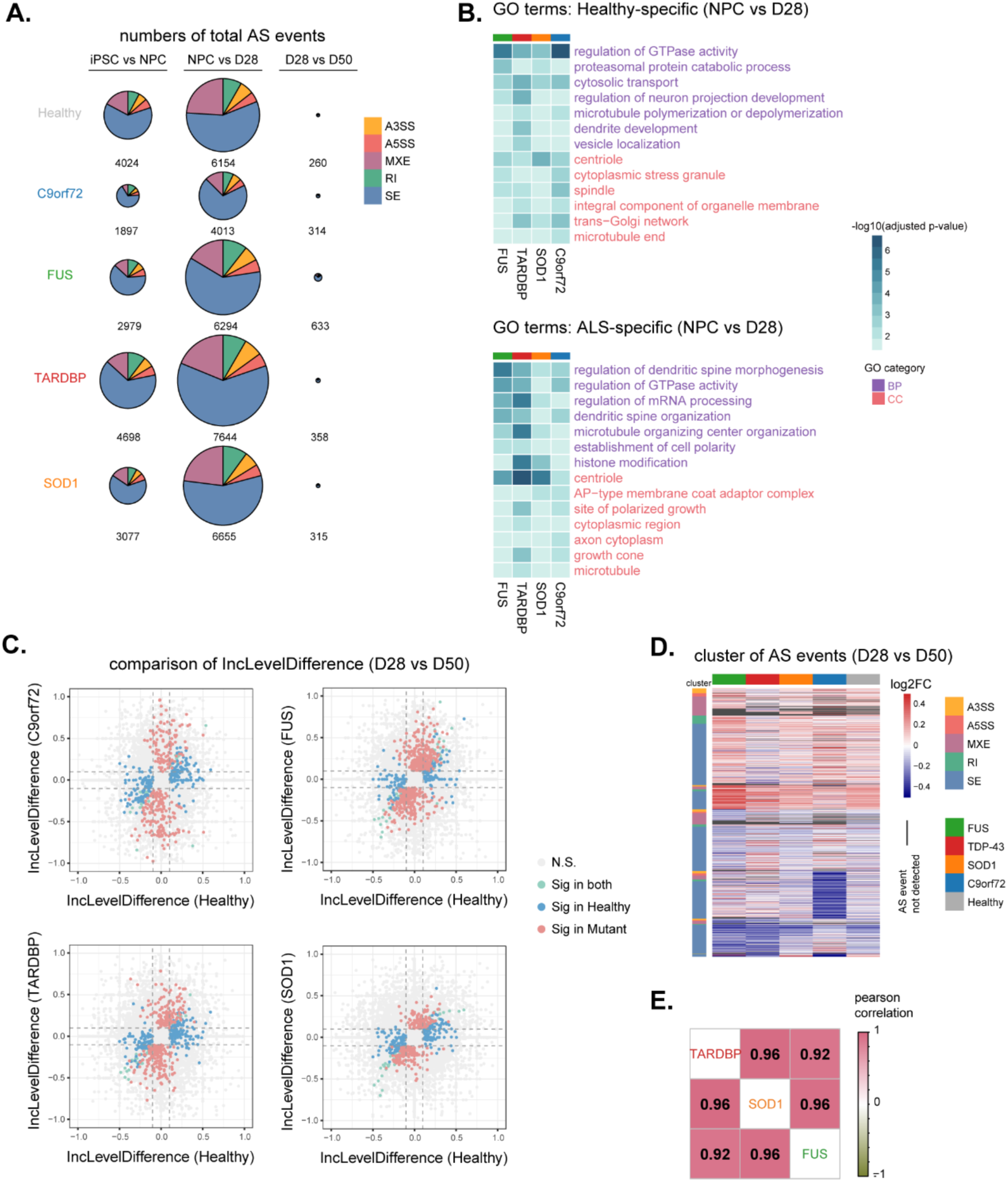
Aberrant splicing events during iMN development and maturation. (A.) Pie charts representing counts of splicing events in each stage of iMN development compared to former stages. Chart areas are proportional to the total numbers of AS types at each stage. (B.) Common GO terms associated with genes only differentially alternative spliced in healthy samples (left) from NPC to day 28 transition compared to corresponding ALS subgroups and vice versa (right). GO biological process (BP) is colored purple, and GO cellular component (CC) is colored red. (C.) Scatter plot of pair-wise comparisons of inclusion level differences of AS events detected in ALS-*C9orf72*, ALS-*FUS*, ALS-*TARDBP*, and ALS-*SOD1* iMNs versus healthy controls. AS events show significant changes in both (green), specific in healthy (blue), and specific in ALS subgroups (red). (D.) Heatmap showing inclusion level differences of AS events significant in at least one iMN subgroup from included (red) to excluded (blue) (1484 events in 1055 genes). AS events that are not detected as significant are depicted with dark grey. (E.) Correlation of inclusion level differences of AS events significant in at least one ALS-*FUS*, ALS-*TARDBP*, and ALS-*SOD1* subgroups.

**Supplementary Fig 5.**
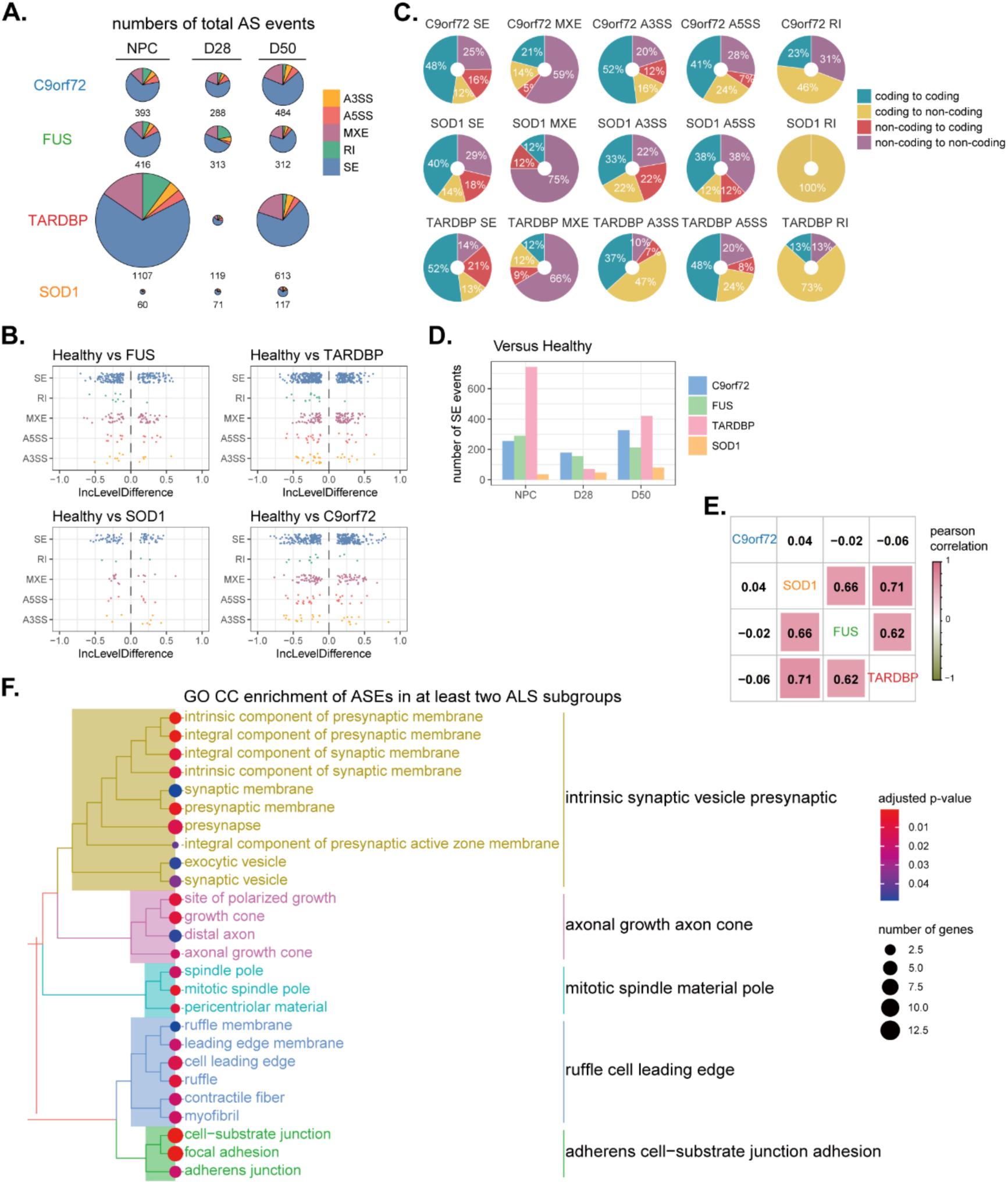
Temporal alternative splicing alterations in ALS iMNs. (A.) Pie charts representing counts of splicing types in ALS subgroups at distinct stages of motor neurogenesis compared to healthy controls. Chart areas are proportional to the total numbers of events at each stage. (B.) Jitter plot displaying distributions of included (IncLevelDifference > 0) and skipped (IncLevelDifference < 0) splicing events of all AS types in ALS iMNs compared to healthy controls. (C.) Pie charts displaying the annotated transcripts’ distribution change on potential protein-coding ability influenced by AS events in ALS-*TARDBP*, ALS-*SOD1*, and ALS-*C9orf72* iMNs on day 50. Coding potentials are calculated by mapping splicing events to transcripts. (D.) Bar graphs illustrating the counts of SE and MXE in ALS subgroups from NPC to D50-iMN in this study. (E.) Heatmap illustrating the correlation of inclusion level differences of AS events significant in at least one of the four ALS subgroups. Larger, darker squares denote a higher Pearson’s correlation coefficient. (F.) Tree plot depicting enriched GO CC terms of genes showing AS significant changes in at least two ALS subgroups.

**Supplementary Fig 6.**
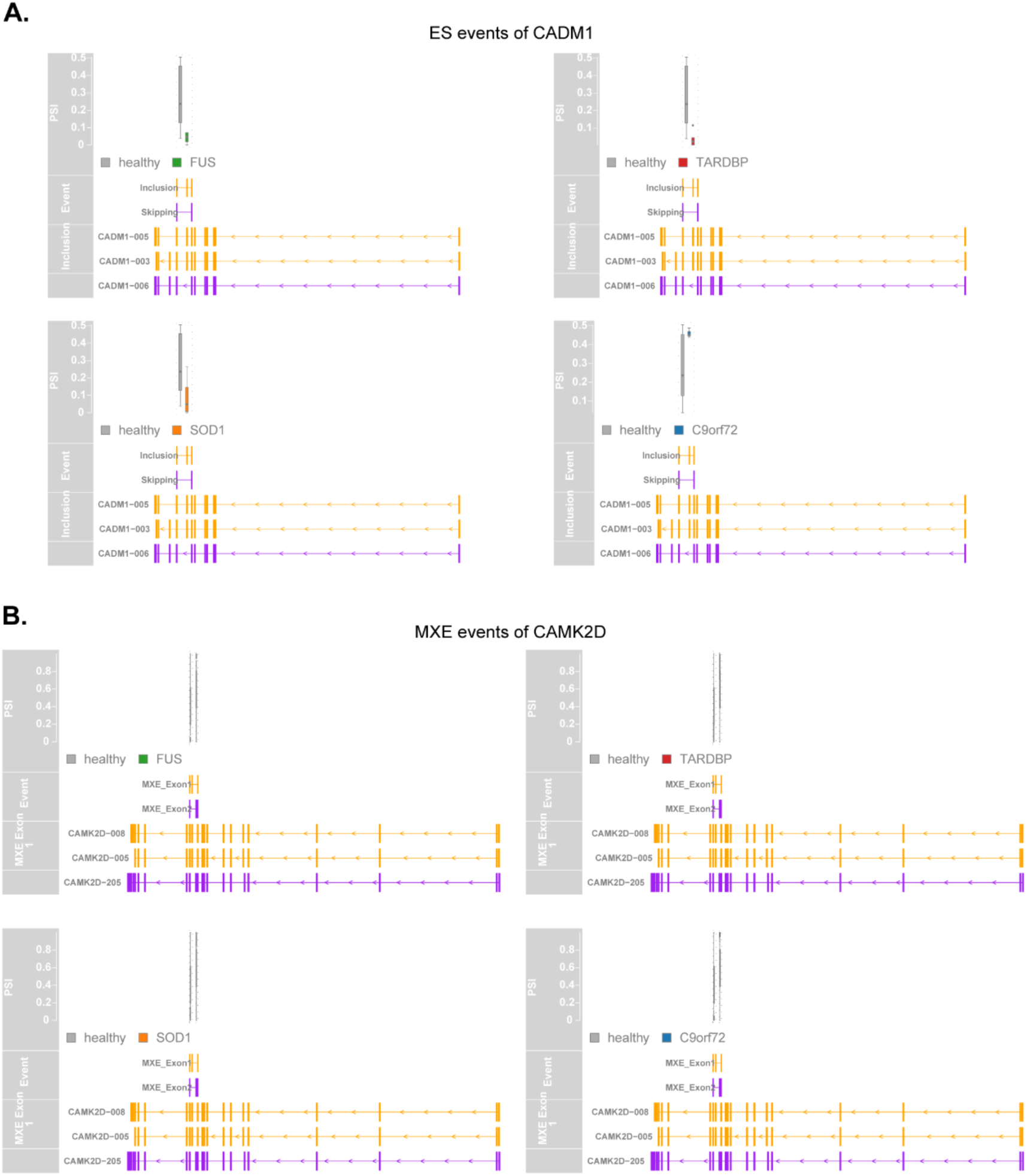
Aberrant AS events of *CADM1* and *CAMK2D* in ALS iMNs on day 50. (A.) Exon skipping events of *CADM1* in ALS iMNs on day 50. (B.) Mutually exclusive splicing events of *CAMK2D* in ALS iMNs on day 50. Boxplots show the difference in inclusion levels (PSI) between healthy and ALS subgroups. The event track showing the AS event with the flanking region, matches the annotated transcript tracks below.

**Supplementary Fig 7.**
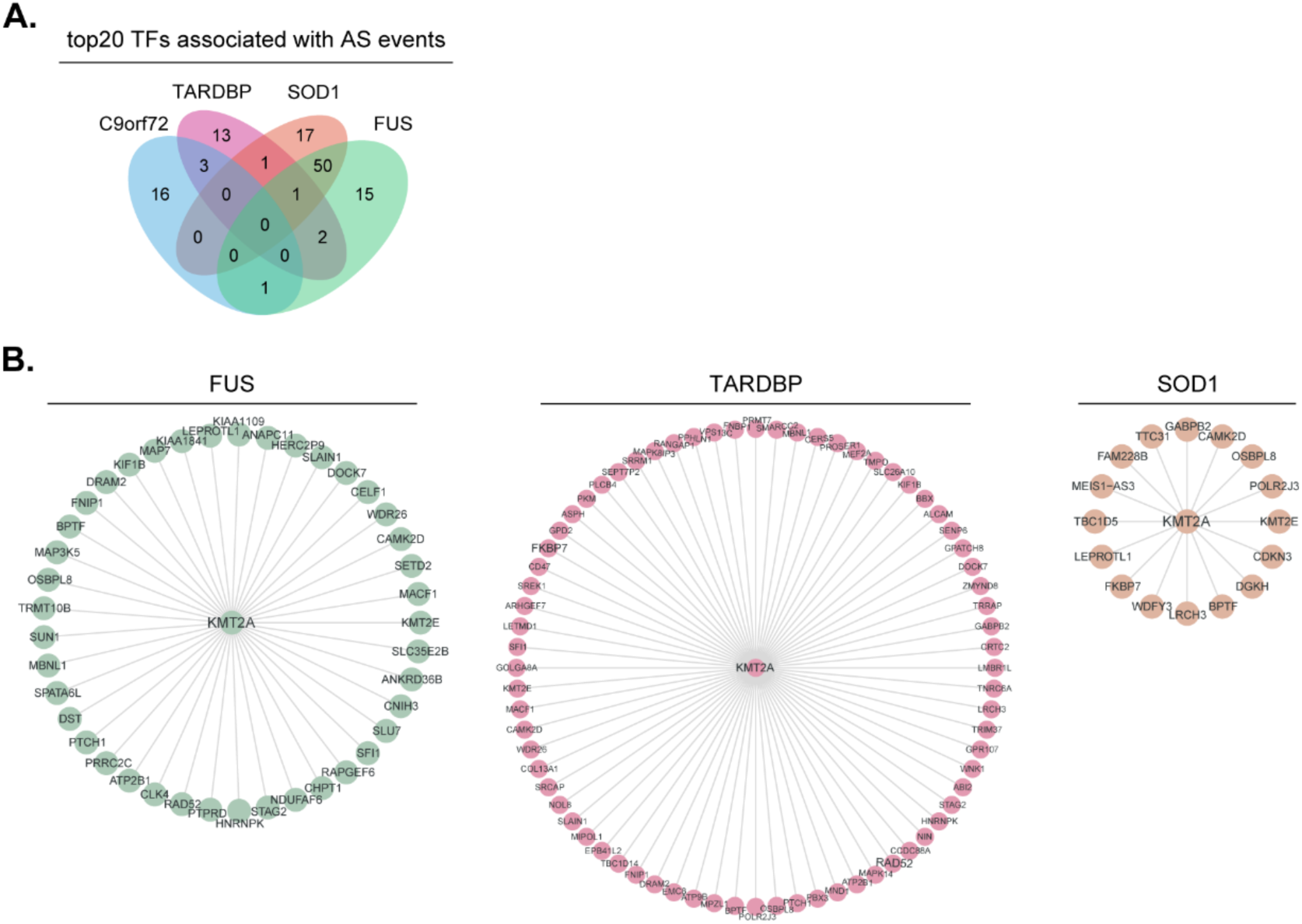
TF regulation of aberrant AS events in ALS iMNs on day 50. (A.) Venn diagram displaying numbers of transcription factors (TFs) associated with AS events that overlap among ALS iMNs. (B.) Networks of key TF KMT2A and AS genes interactions. TF nodes are shown as circles in the center, and their target genes are shown as green (*FUS*), red (*TARDBP*), and orange (*SOD1*) dots. TF-to-AS gene edges are shown as gray lines between nodes.

## Notes

### Competing Interest Statement

The authors have declared no competing interest.

